# A Multifunctional Anchor for Multimodal Expansion Microscopy

**DOI:** 10.1101/2022.06.19.496699

**Authors:** Yi Cui, Gaojie Yang, Daniel R. Goodwin, Ciara H. O’Flanagan, Anubhav Sinha, Chi Zhang, Kristina E. Kitko, Demian Park, Samuel Aparicio, IMAXT Consortium, Edward S. Boyden

## Abstract

*In situ* imaging of biomolecular location with nanoscale resolution enables mapping of the building blocks of life throughout biological systems in normal and disease states. Expansion microscopy (ExM), by physically enlarging specimens in an isotropic fashion, enables nanoimaging on standard light microscopes. Key to ExM is the equipping of different kinds of molecule, with different kinds of anchoring moiety, so they can all be pulled apart by polymer swelling. Here we present a multifunctional anchor, an acrylate epoxide, that enables multiple kinds of molecules (*e.g.,* proteins and RNAs) to be equipped with anchors in a single experimental step. This reagent simplifies ExM protocols and greatly reduces cost (by 2-10 fold for a typical multiplexed ExM experiment) compared to previous strategies for equipping RNAs with anchors. We show that this unified ExM (uniExM) protocol can be used to preserve and visualize RNA transcripts, proteins in biologically relevant ultrastructure, and sets of RNA transcripts in patient-derived xenograft (PDX) cancer tissues, and can support the visualization of other kinds of biomolecular species as well. Thus, uniExM may find many uses in the simple, multimodal nanoscale analysis of cells and tissues.

## Introduction

Nanoscale imaging enables the analysis of biological systems, such as cells and tissues, at the fundamental length scales of biomolecules and biomolecular interactions, but classically has required expensive equipment and advanced skillsets to perform, *e.g.,* via super-resolution microscopy^1^. Recently, expansion microscopy (ExM), which physically magnifies biological specimens through a chemical process, thus enabling nanoimaging on conventional microscopes^2–4^, has become popular, with hundreds of experimental papers and preprints exploring a diversity of biological systems with nanoscale precision^5^. ExM protocols comprise several typical steps: first, one or more molecular anchors are introduced to covalently bond with target biomolecules, or labels bound to those molecules (*e.g.,* endogenous proteins or nucleic acids, or fluorescent probes bound to them); second, acrylate monomers are infused and then polymerized into a swellable polyacrylate hydrogel network that also binds to the anchors; third, the resultant specimen-hydrogel composite is subjected to denaturation or enzymatic digestion to free up inter/intra-molecular connections (*e.g.,* fixative crosslinks), or even to dissolve molecules no longer needed for visualization; finally, the sample is isotropically expanded (typically ∼4.5× in the most commonly used protocols^6^) upon dialysis with an excess of water (**Figure 1A**). ExM has been successfully demonstrated in a wide range of sample types and given rise to a number of variants tackling specialized purposes, *e.g.,* higher magnification factors, adaptation to human tissues, decrowding of molecules for better access by labels, and multiplexed molecular imaging^7–24^, to name a few, which have been used to study a diversity of topics in virology, molecular biology, neuroscience, cancer biology, and other fields within biology and medicine.

**Figure 1.**
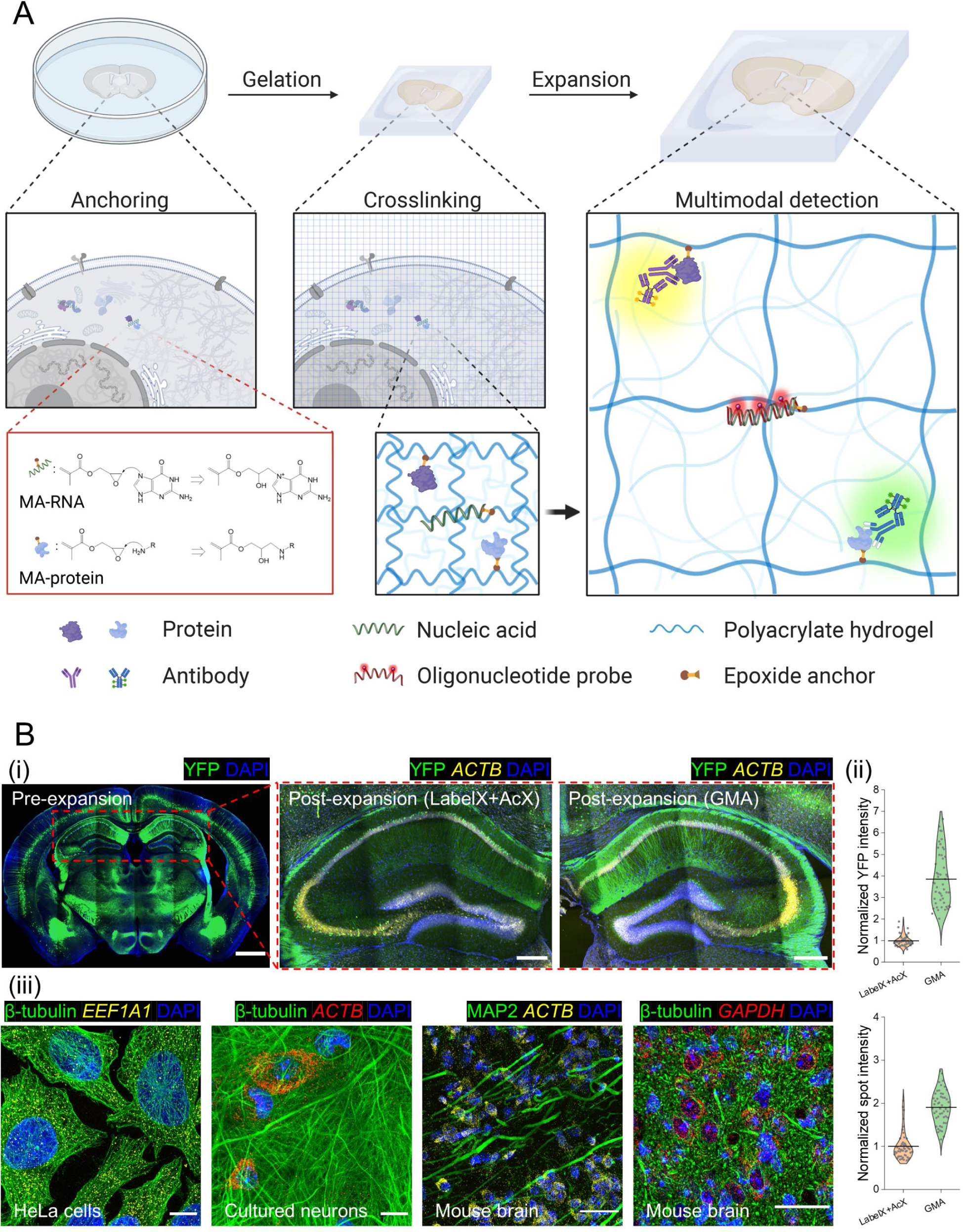
An epoxide anchor enables expansion of multiple kinds of biomolecule away from each other (uniExM). **(A)** In a standard ExM experiment, target biomolecules (*e.g.,* proteins and nucleic acids, or labels attached to them) in a biological specimen are covalently bound to anchoring molecules bearing vinyl groups (here, an epoxide such as glycidyl methacrylate (GMA), turning biomolecules to methacrylate (MA) forms, *e.g.,* MA-RNA and MA-protein) that can be crosslinked to a swellable polyacrylate hydrogel synthesized throughout the specimen (“Gelation”). After tissue softening with denaturation and/or proteolysis, the sample can be isotropically expanded upon dialysis with low osmolarity solutions (*e.g.,* distilled water), during which the anchored biomolecules are pulled apart (“Expansion”). Target-specific detection can be performed (*e.g.,* antibody staining for proteins and oligonucleotide probe hybridization for RNAs) to realize nanoimaging on conventional microscopes. **(B)** Using the epoxide anchor GMA, simultaneous detection of nucleic acids and proteins was demonstrated across different sample types. In panel **(i)**, a 50 µm thick coronal section of mouse brain tissue expressing Thy1-YFP was cut in half, one part anchored with 0.01% (w/v) LabelX plus 0.005% (w/v) AcX and the other with 0.1% (w/v) GMA. The samples were then subjected to gelation and proK based digestion. HCR-FISH targeting *ACTB* mRNAs was performed post-expansion and the FISH signals were quantified together with the retained YFP signals. Scale bars (in pre-expansion units): 1000 µm (whole brain); 250 µm (zoomed-in hippocampal view). Linear expansion factor: 4.1. In panel **(ii)**, mean intensities for YFP within cells and HCR-FISH spots in each image were quantified and compared between the LabelX/AcX and GMA processed tissues (data distribution shown in violin plots, with raw data points presented, and mean values highlighted with solid lines; n = 50 images from 3 different slices, 2 mouse brains; two- sample *t*-test was performed with *p* < 10^-20^ for both YFP and HCR-FISH signals). In panel **(iii)**, multimodal detection of proteins and RNAs with uniExM was demonstrated. Antibody staining for proteins was performed pre-expansion and the samples were anchored with 0.04% (w/v) GMA (for HeLa cells and cultured neurons) or 0.1% (w/v) GMA (for mouse brain slices). After gelation and expansion, HCR-FISH targeting specific RNA targets was performed post-expansion. In the figure captions, capitalized italic fonts represent the RNA target names (*ACTB, EEF1A1, GAPDH*). The colors in each image correspond to the following fluorescent dyes: blue – DAPI; green – Alexa488; yellow – Alexa546; red – Alexa647. Scale bars (in pre-expansion units): 20 µm. Linear expansion factors: 4.4 for HeLa cells and cultured neurons, 4.1 for mouse brain tissues. All images are shown as maximum z-projection of image stacks (10-15 µm z-depth for cultured HeLa cells and neurons; 50 µm for mouse brain tissues).

Although most papers performing ExM investigate a single kind of biomolecule, *e.g.,* proteins, an increasing number of studies are seeking to investigate multiple kinds of molecule, *e.g.,* both proteins and RNAs^4, 21, 25, 26^. Proteins and RNAs to date have required different anchors – proteins have been bound by molecules that connect amines to a vinyl group (*e.g.,* through the molecule AcX)^3, 7, 8^, whereas RNAs have been bound by molecules that alkylate guanines (*e.g.,* through the molecule LabelX)^4, 25^ (schematics in **Supplementary Figure 1**). The RNA anchors are made by mixing off-the-shelf chemicals overnight; the RNA anchors are sometimes applied to the specimen in a separate step from the protein anchors, adding time and complexity to the procedure. The need for in-house synthesis adds a potential uncertainty to the overall process, since individuals may not conduct the reaction under identical conditions in all groups, resulting in non-controlled final yield. The cost of the RNA-binding reagents is also high – LabelX and MelphaX (AcX reacted with Label-IT amine and Melphalan, respectively), the two molecules used to date, cost $7,500 and $70 per mg, respectively, as of May 2022, resulting in overall experimental costs of $180 and $40, respectively, for a typical sample (*e.g.,* a 50-100 µm thick full-width mouse brain slice.

Here we introduce, and validate, an epoxide-based anchoring strategy that enables multiple molecular species, such as proteins and RNAs, to be anchored to the hydrogel in a single step, in an inexpensive fashion, without requiring any end-user synthesis. We demonstrated the versatility of this unified ExM (uniExM) protocol in the analysis of proteins and RNAs, as well as in multiplexed settings such as that of expansion sequencing (ExSeq)^21^, suggesting the utility of epoxide anchoring in a diversity of high-resolution spatial biology studies.

## Results

### Epoxides as multifunctional anchors

We reasoned that an ideal multifunctional anchor for ExM should be chemically active, mechanistically predictable, and universally accessible. Epoxides fit this bill, and are already ubiquitous in daily life (*e.g.,* in epoxy adhesives) and well understood. The ring-opening process of epoxides is a nucleophilic substitution reaction and could follow two pathways: an SN1-like reaction under acidic conditions or an SN2 reaction under basic conditions, making the anchoring reaction pH sensitive^27^. Acidic solutions are able to protonate epoxides and open the high-tension three-atom ring directly, resulting in rapid conjugation with weak nucleophiles such as water and alcohol.^28^ However, acids could also protonate nucleophiles on the biomolecules or labels to be anchored. Hence, we chose a slightly basic system, well within the range of standard specimen treatments (pH = 8.5, buffered by 100 mM sodium bicarbonate NaHCO3), for epoxides to react with nucleophiles of biological importance, including but not limited to cysteine, histidine, lysine, glutamic acid, tyrosine, and guanine (schematics in **Supplementary Figure 2**)^12, 29, 30^. We chose for this paper to investigate glycidyl methacrylate (GMA), which contains an epoxide and a vinyl group, as an anchoring moiety. A high-level comparison of LabelX, MelphaX, and GMA with regards to key properties in application is summarized in **Supplementary Table 1**.

We confirmed that uniExM is compatible with both a popular protein retention form of ExM (proExM) and an RNA retention form of ExM that supports *in situ* hybridization (ExFISH) (examples in **Supplementary Figure 3** and **Figure 1Bi**). Specifically, 0.04% (w/v) GMA, as compared to an established method using 0.01% (w/v) LabelX plus an additional 0.005% (w/v) AcX^4^, fared well, retaining native yellow fluorescent protein (YFP) intensity ∼4 times stronger (**Figure 1B, ii)**. The single-spot intensity of HCR-FISH was 2 times stronger in GMA treated tissues (**Figure 1B, ii**), reminiscent of the improvement yielded by MelphaX over LabelX^25^. Both RNAs and proteins could be retained and visualized in the same specimen, after GMA anchoring, for both cultured cells and intact tissues (**Figure 1B, iii**); hence, we termed this simplified protocol unified expansion microscopy (uniExM).

### Characterization of uniExM distortion and yield

Perhaps because only the anchoring step was changed, without alteration of the gelation, softening, and expansion steps, the rest of the expansion process proceeded smoothly, without requiring further refinement. Not surprisingly, uniExM supported high-resolution imaging of fine structures, including microtubules in cultured HeLa cells (**Figure 2A**). The expansion factor was found to be 4.2-4.4 using 0.04% GMA (**Figure 2A, iv**) and the distortion incurred by expansion was similar to previously published results, a few percent over a field of view of a few tens of microns (benchmarked against Nikon SoRa super-resolution microscopy) (**Figure 2A, v**). In addition, the morphology and geometry of the nucleus was reliably maintained after expansion (**Supplementary Figure 4**).

**Figure 2.**
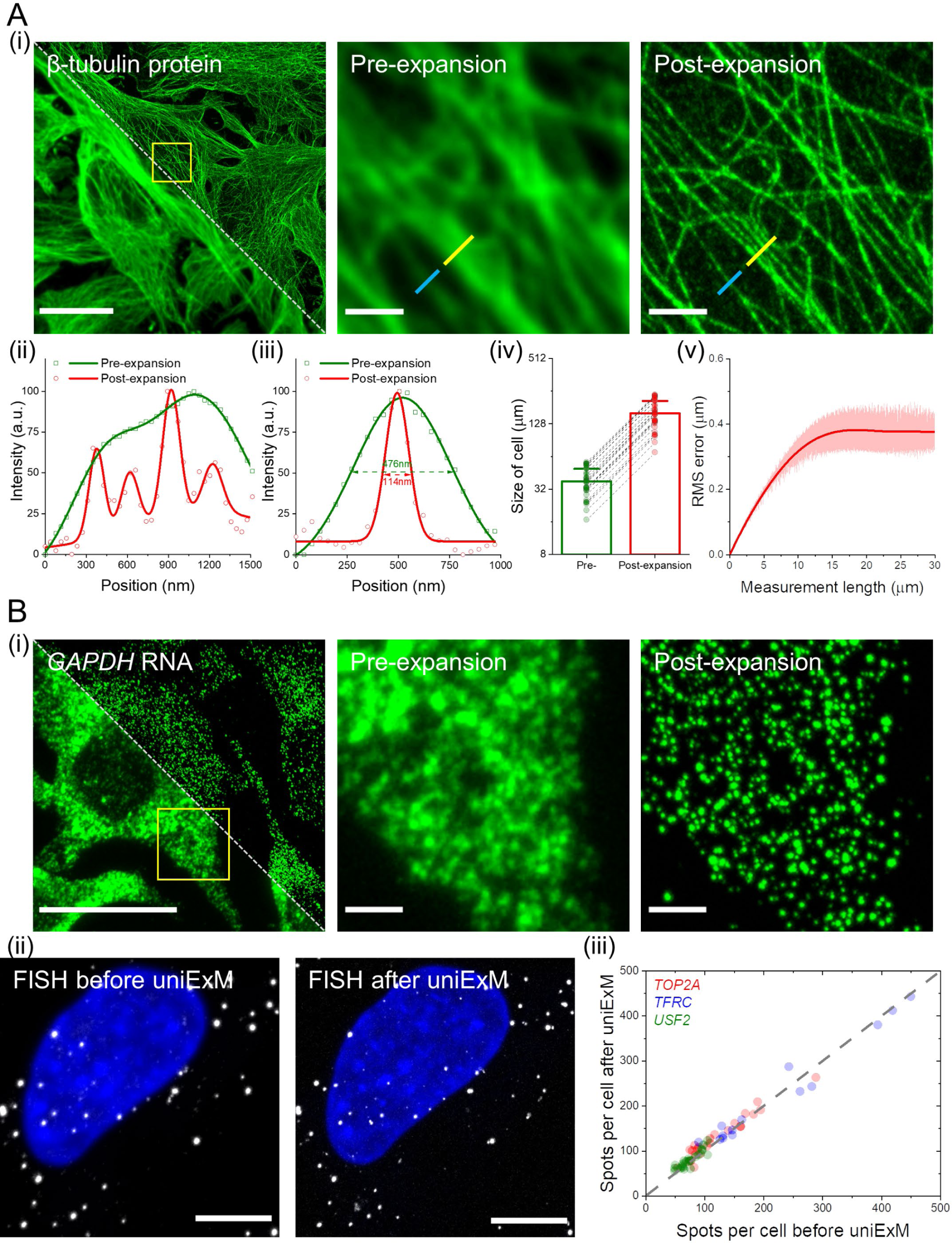
Characterization of GMA-based uniExM for protein and RNA retention. **(A)** uniExM improves imaging resolution and achieves homogenous expansion. **(i)** Representative images of HeLa cells stained with β-tubulin antibody (pre-expansion staining) are shown. Upon expansion, resolution improvement, expansion factor and distortion were evaluated. Left image: pre-expansion (lower left half) and post-expansion (upper right half) fields of view of the same specimen, where the white diagonal dashed line delineates the boundary between the two images. Middle and right images: zoomed-in view of the region highlighted by the yellow square in the left image. Scale bars (in pre-expansion units): 20 µm (left image), 2 µm (middle and right images). Panels **(ii)** and **(iii)** plot the cross-section intensity profiles along the yellow and blue lines, respectively, in the zoomed-in images of panel **(i)**. The raw intensity values (shapes) were fitted with multi-peak Gaussian functions (solid lines). The values presented in panel **(iii)** are FWHM (full width at half maximum) of the fitted Gaussian functions. In panel **(iv)**, long axes of the same cell were measured before and after expansion to calculate the expansion factor. The linear expansion factor was determined to be 4.2 in this demonstration (n = 30 cells from 2 different batches of culture; mean + standard deviation was presented in bar chart with raw measurements shown as individual points). In panel **(v)**, RMS length measurement error was quantified by benchmarking post-expansion confocal images against pre-expansion super-resolution SoRa images of microtubule staining in HeLa cells (red line, mean value; shaded area, standard deviation; n = 5 samples). **(B)** uniExM for RNA detection and quantification. **(i)** GMA-based expansion helps de-crowd densely packed mRNAs and better resolve single transcripts of the highly expressed *GAPDH* gene in HeLa cells. Left image: pre-expansion (lower left half) and post-expansion (upper right half) images of the same specimen, where the white diagonal dashed line delineates the boundary between these two images. Middle and right images: zoomed-in view of the region highlighted by the yellow square in the left image. Scale bars (in pre-expansion units): 20 µm (left image), 2 µm (middle and right images). (ii) GMA-based uniExM effectively preserves RNA information during the expansion process. HCR-FISH targeting specific genes was performed before and after expansion. Left image: a representative image of HCR-FISH for the *USF2* gene in HeLa cells. Number of transcripts per cell was counted, and then FISH probes were stripped off with concentrated formamide and heating. Right image: The same sample was subjected to uniExM, after which HCR-FISH targeting the same gene was performed and quantified. Scale bars (in pre-expansion units): 5 µm. **(iii)** Three genes – *TOP2A, TFRC and USF2* (with the expression level ranging from ∼50 to ∼500 transcripts per cell) – were chosen to evaluate the RNA anchoring efficiency by GMA in uniExM. Spots/transcripts per cell counted before and after GMA anchoring were fit by linear regression, with an R-squared value of 0.9736, indicating nearly 100% RNA retention (each point in the scatter plot represents one measurement from a single cell; n = 60 cells collected from 3 culture batches).

As with ExFISH before it, uniExM is able to decrowd densely packed RNA molecules for better detection (**Figure 2B, i**). To gauge how uniExM facilitates retention of RNA molecules, we systematically varied GMA concentration, reaction pH, and temperature, evaluating the retention of three highly expressed genes (*GAPDH, EEF1A1, ACTB*) (**Supplementary Figure 5**). The optimal reaction condition was determined to be

0.04% GMA for cultured cells at pH 8.5 (100 mM NaHCO3), at room temperature. For tissue samples we used 0.1% GMA to ensure sufficient anchoring of multiple types of biomolecules, and found in practice that increased temperature (*e.g.,* from room temperature to 37°C) could facilitate the diffusion and anchoring efficiency of epoxide, consistent with previous reports^4, 6^. Moreover, we demonstrated that the anchoring reaction could be efficiently controlled by varying temperature and pH; at 4°C and neutral pH, the anchoring efficiency for RNA can be suppressed by more than 50% after 12 h incubation (termed reaction “Off” condition), while it can be rapidly recovered to full efficacy by an additional 3 h incubation at 25°C and pH 8.5 (termed reaction “On” condition) (**Supplementary Figure 5D**).

We quantified three moderately expressed genes (*TOP2A, TFRC, USF2*) with hybridization chain reaction (HCR)-FISH before vs. after GMA anchoring. uniExM preserved RNA molecules with ∼100% retention efficiency (**Figure 2B, ii-iii**). As another comparison, GMA was equal to, or perhaps even slightly better, than LabelX in preservation of highly expressed mRNA targets (**Supplementary Figure 6**). Thus, both in terms of supporting even expansion, and high yield of target, GMA is an excellent unified anchor for both proteins and RNAs.

### uniExM for preservation of protein content and ultrastructure

We next explored applications of uniExM in different biological contexts. We chose βII- spectrin in neurons to image, since its periodic distribution in axons was discovered via super-resolution imaging^31, 32^. In cultured mouse hippocampal neurons, the periodic distribution of βII-spectrin was prevalently observed in axons (**Figure 3A**; more examples in **Supplementary Figure 7**). The distance between two adjacent βII-spectrin spots was found to be ∼190 nm (**Figure 3B**), as reported earlier^31^, this periodicity was not apparent in pre-expansion samples. The periodic distribution of βII-spectrin can be additionally visualized with an autocorrelation analysis (**Figure 3C**), which we performed with a 7× expansion protocol based on the TREx protocol^24^. Finally, autocorrelation analyses from four independent samples were performed, yielding calculated periodicity values (193 ± 15 nm for 4× expanded cells, 187 ± 10 nm for 7× expanded cells, values reported as mean ± standard deviation) consistent with results previously obtained by super-resolution STORM or STED imaging (**Figure 3D**)^33, 34^.

**Figure 3.**
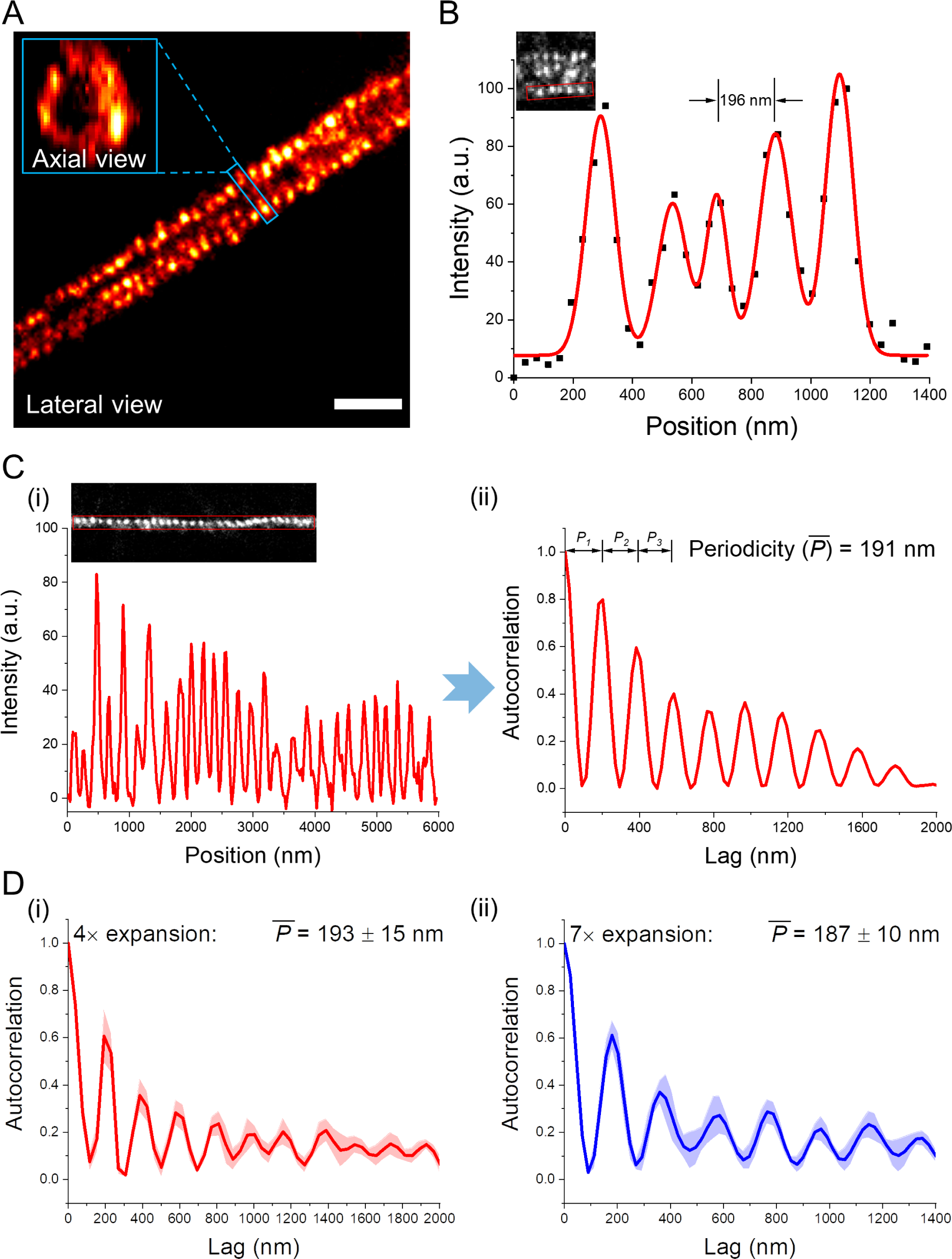
Preservation of protein ultrastructural organization by GMA in uniExM. **(A)** Antibody staining against βII-spectrin in mouse hippocampal neurons was performed with 4× expansion (linear expansion factor ∼4.2). One segment of axon showing periodic, punctate signals is presented. In the zoomed-in inset, the axial view of βII-spectrin is shown, displaying a ring structure. Scale bar (in pre-expansion units): 1 µm. **(B)**The intensity profile of βII-spectrin clusters along an axon segment (within the red rectangle of the inset image) was plotted and fitted with multi-peak Gaussian functions (black squares: raw data points; red line: fitting function). A consistent distance around 190 nm between two adjacent peaks was noted. **(C)** Antibody staining against βII-spectrin in mouse hippocampal neurons was performed with a 7× expansion protocol modified from the TREx protocol (linear expansion factor ∼7.3). **(i)** The intensity profile of over 10 βII-spectrin clusters along an axon segment (within the red rectangle of the inset image) was plotted. **(ii)** Autocorrelation analysis was performed to calculate the periodicity of βII-spectrin clusters across space (*i.e.,* the similarity between signals as a function of the spatial position lag between them). Based on each fitted autocorrelation function, the first three inter-peak distance values (denoted as *P_1_, P_2_* and *P_3_*) were extracted to calculate the mean periodicity *P̅* . **(D)** Summary of periodicity values obtained under different expansion factors. **(i)** Mean autocorrelation function of periodicity analysis under 4× expansion. **(ii)** Mean autocorrelation function of periodicity analysis under 7× expansion (solid lines, mean; shaded areas, standard error of mean. n = 40 measurements from 4 samples, 2 culture batches).

Polyepoxides, which can crosslink multiple biomolecules to each other, and even multiple parts of a biomolecule to each other, have been shown to help proteins such as fluorescent proteins retain function in tissue specimens under harsh conditions, such as high heating^12^. We combined GMA with a polyepoxide, trimethylolpropane triglycidyl ether (TMPTE), to assess the performance of this cocktail in preservation of fluorescent protein function after heat-based tissue softening. We tested two tissue processing protocols requiring heating above 50°C: the SDS-based sample denaturation (95°C for 1h)^19^ and the proK-based enzymatic digestion (60°C for 2h)^35^. Compared with LabelX plus AcX, GMA plus TMPTE showed better retention of fluorescence signals for paraformaldehyde (PFA)-perfused, fresh frozen Thy1-YFP mouse brain slices (∼440% signal improvement for SDS-based denaturation; 250% signal improvement for proK- based digestion; **Supplementary Figure 8**).

### uniExM for *in situ* sequencing of RNA

Expansion microscopy greatly facilitates *in situ* sequencing, enabling multiplexed RNA analysis with high spatial precision; we named the optimized combination expansion sequencing (ExSeq)^21^. In untargeted ExSeq, linear probes containing randomized octamer sequences are hybridized to RNA targets, followed by reverse transcription of adjacent RNA sequence information into cDNA form. cDNAs are then circularized and undergo rolling circle amplification (RCA) (schematized in **Supplementary Figure 9**). Then, in each round of sequencing readout, the bases being added are imaged through fluorescence microscopy (example in **Supplementary Figure 10A**). In addition to untargeted ExSeq, one can perform targeted ExSeq against sets of specific RNAs (**Figure 4A**), by bringing in padlock probes that hybridize to targets, and then sequencing barcodes found on the padlock probes (schematized in **Supplementary Figure 9**; example in **Supplementary Figure 10B**). Comparing amplicons (containing amplified barcode sequences) from targeted ExSeq using GMA, to spots seen with classical HCR-FISH, showed excellent agreement in spot counts (**Figure 4B, i**). The sequencing, conveniently, can be done with standard Illumina MiSeq sequencing-by-synthesis (SBS) reagents (**Figure 4B, ii**). We performed sequencing of barcodes on padlock probes targeting *ACTB* mRNAs to test the stability of uniExM-based ExSeq across multiple rounds of sequencing in mouse brain tissues. The padlock probes used contained repetitive “T” bases in their barcode region, so the amplicons should emit the same fluorescence signal in each round of sequencing. The amplicons were first examined with a universal detection probe to generate a reference image, then three consecutive rounds of sequencing were performed (**Supplementary Figure 10C**).

**Figure 4.**
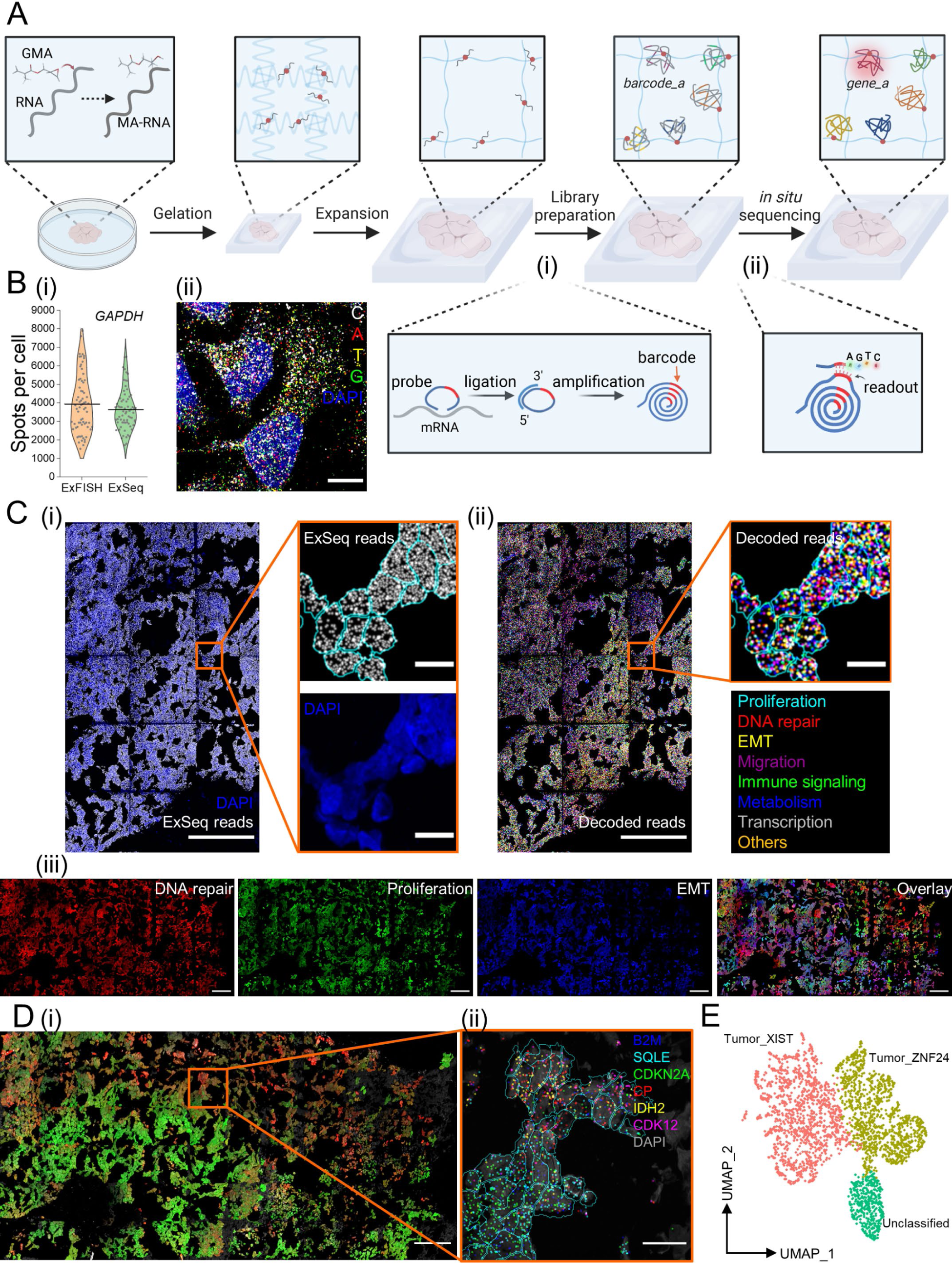
uniExM-supported *in situ* RNA sequencing (ExSeq). **(A)** Schematic of the workflow for targeted ExSeq (tExSeq): target RNA molecules are reacted with GMA to acquire methacrylate groups (termed MA-RNA) and anchored to the expandable hydrogel. Padlock probes are then introduced to hybridize with the target RNAs in the expanded biological sample. Upon successful hybridization, the sequence of a target mRNA serves as a “splint” for PBCV-1 enzyme-mediated ligation of the bound padlock, as shown in the zoomed-in panel **(i)**. Afterwards, rolling circle amplification (RCA) is applied to amplify the ligated probes that harbor barcodes specific to targets. Finally, the barcodes are read out by *in situ* sequencing chemistry, as shown in the zoomed-in panel **(ii)**. The full readout of a specific barcode can then be used to reveal the gene identity (*e.g.,* from barcode_a to gene_a) together with its location information. In such way, multiple gene targets can be decoded (represented as differentially colored amplicons). **(B)** Validation of ExSeq enzymatics and sequencing-by-synthesis (SBS) chemistry in samples processed with the uniExM procedure. **(i)** *GAPDH* in HeLa cells was chosen to undergo HCR-FISH or tExSeq. The numbers of detected signal spots per cell were quantitatively compared. No statistically significant difference was observed between the two methods. (Data shown as violin plots, with raw data points presented, and mean values highlighted with solid lines; n = 70 cells from 4 samples, 2 culture batches; two-sample *t-*test was performed with *p >* 0.1) **(ii)** A fluorescence image showing raw ExSeq signals from all four base channels in HeLa cells undergoing targeted ExSeq. Scale bars (in pre-expansion units): 20 µm. **(C)** Demonstration of ExSeq applying an 87-gene panel in GMA-anchored SA501 PDX breast cancer tissue. **(i)** Overview of the raw ExSeq reads (gray spots) in the tissue. DAPI staining for nuclei was used for cell segmentation and reads assignment (shown in the zoomed-in images). **(ii)** The raw ExSeq reads were decoded and colored based on 8 distinct gene function groups (full list in **Supplementary Table 5**). Scale bars (in pre-expansion units): 100 µm (for stitched overview images), 10 µm (for zoomed-in images). **(iii)** Gene maps of 3 selected function groups – DNA repair, proliferation and epithelial- mesenchymal transition (EMT), were used to help visualize heterogeneity of cell status in the whole tissue. The decoded transcripts of genes belonging to each functional group were summed and their ratio to total transcript counts were assigned to the “R”, “G”, “B” color channels of the image, respectively. During the color assigning process, a scaling factor of 3.33 (for DNA repair and proliferation) or 2.5 (for EMT) was introduced; that is, if the EMT group of genes was 40% of the total transcripts in one cell, its assigned “B” channel was given the maximum color intensity (40% X 2.5 = 100%). Then, the three individual channels were combined to make a composite image (right). Scale bars (in pre-expansion units): 100 µm. Linear expansion factor: 3.2 (the expanded gel was re-embedded before sequencing). **(D)** Unsupervised principal component analysis (PCA) identified two primary gene groups for cell classification. **(i)** Using these two PCA gene groups, a distribution of different cancer cells was revealed. In this presented image, the summed transcripts of each PCA group normalized to the total transcript count for a given cell were assigned to the “R” channel and “G” channel, respectively (scaling factor: 3.33). Then the two images were overlaid to make a composite. Scale bar (in pre-expansion units): 100 µm. **(ii)** In the zoomed-in region where differentially colored cells co-exist, 6 marker genes are plotted; their distribution varies across cells in the subregion. Scale bar (in pre-expansion units): 10 µm. **(E)** Uniform manifold approximation and projection (UMAP) representation of the cell typing results using bulk RNA-seq identified marker genes in the SA501 cancer model. According to the panel design, the 87 gene list could differentiate two primary cancer cell clones that are successfully annotated on UMAP – “Tumor_XIST” and “Tumor_ZNF24”, named after their feature genes. A small group of cells are marked as “Unclassified”, likely attributed to non-cancer interstitial or other cells.

Amplicons were consistently detected in each round (with an average detection rate of ∼96.5% across 20 fields of view from 2 different mouse brains), indicating that uniExM- based ExSeq is compatible with the full SBS chemistry cycle (*i.e.,* elongation, detection and cleavage steps). Moreover, Thy1-YFP positive structures in the mouse brain tissues (*e.g.,* dendrites and spines) were consistently detected after ExSeq library preparation procedure (as seen by comparing images from the YFP channel before vs. after library preparation), helpful for analyzing transcriptomics in the context of morphological compartments like synapses and axons.

We performed 7-round SBS to profile 87 genes known to classify tumor cells into subclasses in SA501 breast cancer PDX tissues^36^ (the original LabelX-based ExSeq was demonstrated for 4 rounds of SBS). Importantly, uniExM significantly reduces the cost of ExSeq, as the anchoring reagent contributes to more than half of the entire cost of the original protocol (see **Supplementary Table 1** for detail). In a 1.24 × 0.62 mm^2^ region of interest, 793,535 raw reads (colored by function annotations) and 3,339 cells in total were detected (**Figure 4C, ii**; **Supplementary Figure 10D**). Many kinds of analysis are possible: in **Figure 4C, iii**, transcripts from three functional groups were presented as 8-bit “RGB” images where the color intensity reflected the overall expression level of the members of that group in each individual cell. Principal component analysis (PCA), applied to all the cells of the sample, revealed two distinct cell groups distinguished by 30 out of the 87 genes (**Supplementary Figure 10E**).

Based on the expression levels of these two PCA-generated gene sets, we assigned a color code to each individual cell (**Figure 4D, i**). The summed counts of each PCA group-specific gene set (15 genes), normalized to the total count in each individual cell, were then plotted as the red and green channels of the “RGB” image, respectively. The composite image shows a spatially varying distribution of the colored cells in space, explicitly delineating two tumor cell populations. In a zoomed-in region where cells of both groups were present, transcripts from distinct PCA lists were detected in different cells (**Figure 4D, ii**). As a second method of analysis, we ran a dimensionality reduction algorithm using the clone-specific marker genes with significant differential expression from the RNA-seq data of the same PDX model^37^ and presented the results with UMAP, where two tumor cell clones (denoted as “Tumor_XIST” and “Tumor_ZNF24”) and one group of “Unclassified” cells were identified (**Figure 4E**). These two classified groups of cells well correspond to the bright-green and bright-red colored cells in **Figure 4D, i**, respectively. In detail, comparing the unsupervised and supervised results, 85.1% of tumor clone cells annotated by the supervised method were classified as the same clone by unsupervised method, whereas 82.7% of cells classified by the unsupervised method were cross-verified by the supervised method.

### uniExM for multimodal detection beyond proteins and nucleic acids

We examined the compatibility of uniExM with commercially available lipid stains and markers of carbohydrates. We selected three lipid stains – octadecyl rhodamine B chloride (R18), FM 1-43FX (FM) and BODIPY FL C12 (BODIPY), all of which showed signals in lipid-rich or membranous structures (**Supplementary Figure 11A-B**) . Using Alexa647 tagged wheat germ agglutinin (WGA), which binds *N*- acetylglucosamine (GlcNAc), we were able to see specific signals in nuclear membranes of cultured HeLa cells and blood vessels of mouse brain tissues (**Supplementary Figures 11C and 12**). Such stains could be used together, in the same specimen, along with antibody stains and FISH probes, supporting multimodal imaging of many kinds of biomolecule in the same cell or tissue slice (**Supplementary Figure 12**).

## Discussion

Here we show that a single anchoring molecule can support the expansion of both proteins and RNAs away from each other in expansion microscopy, reducing the complexity and cost compared to earlier anchoring strategies, which required different anchors for different kinds of biomolecules. We used an epoxide, an inexpensive and highly reactive moiety capable of binding to many kinds of biomolecule, that contained a vinyl group capable of participating in polymerization. We found this molecule, GMA, to enable good retention of proteins and RNAs, and to support labeling for visualization of proteins, RNAs, and other biomolecules.

Past anchoring strategies for ExM have used different anchors for different kinds of biomolecules, *e.g.,* using an aldehyde or NHS ester to bind amines on proteins, or an alkylating reagent to bind guanine on RNA. The latter strategy requires end users to mix multiple chemicals overnight to create the anchor, and some protocols administer different anchors in separate steps. Here, no end user synthesis is needed, and only a single step is needed to add the universal anchor GMA.

We demonstrated the utility of this unified ExM, or uniExM, protocol in cells and tissues. Perhaps because of the rest of the protocol is unaltered compared to previous ExM protocols, we saw no differences in the performance of the rest of the protocol (*e.g.,* distortion, resolution). uniExM was able to support not just classical ExM protocols like antibody detection of proteins and FISH detection of RNAs, but also our recently published ExSeq protocol for multiplexed visualization of RNAs. In short, uniExM will find utility in a variety of contexts in biology.

## Methods and Materials

### Cell culture and mouse brain tissues

HeLa cells were cultured in DMEM medium supplemented with 10% FBS and 1% penicillin-streptomycin antibiotics. Once the cells reached 70-80% confluence, they were fixed with either 4% paraformaldehyde (PFA) or 3% PFA/0.1% glutaraldehyde (GA, for better preservation of intracellular fine structures including β-tubulin and ΒII- spectrin in **Figures 1B iii, 2A, 3**, **Supplementary Figures 7, 12B**), followed by residual aldehyde quenching with 0.1% sodium borohydride and 100 mM glycine. For RNA detection (ExFISH and ExSeq), samples were permeabilized and stored with 70% ethanol at 4°C overnight or up to 4 weeks.

All procedures involving animals were in accordance with the US National Institutes of Health Guide for the Care and Use of Laboratory Animals and approved by the Massachusetts Institute of Technology Committee on Animal Care. Primary neurons were dissected from brains isolated from euthanized newborn Swiss Webster mice and about 1,000 hippocampal neurons were seeded onto a 12 mm #1.5 coverslip. The neurons were cultured with MEM medium containing 33 mM glucose, 0.01% transferrin, 10 mM HEPES, 2 mM Glutagro, 0.13% insulin, 2% B27 supplement, and 7.5% heat-inactivated FBS at 37°C in a humid incubator supplemented with 5% CO2 for 2 weeks and then fixed for subsequent uses.

For mouse brain tissues, seven-week old mice were terminally anesthetized with isoflurane, followed by transcardial perfusion with PBS and ice cold 4% PFA. Then the brain was dissected out and placed in 4% PFA for 12-16 hours. 50 μm slices were prepared on a vibratome (Leica VT1000s) and then stored at 4°C in PBS (for lipid co-detection in **Supplementary Figure 12**) or 70% ethanol until use.

Patient-derived xenografts (PDX) Patient tumor samples were acquired according to procedures approved by the Ethics Committees at the University of British Columbia. Breast cancer patients undergoing diagnostic biopsy or surgery were recruited and samples collected under protocols H06- 00289 (BCCA-TTR-BREAST), H11-01887 (Neoadjuvant Xenograft Study), H18-01113 (Large scale genomic analysis of human tumors) or H20-00170 (Linking clonal genomes to tumor evolution and therapeutics). Tumor fragments were finely chopped and mechanically disaggregated for one minute using a Stomacher 80 Biomaster (Seward Limited, Worthing UK) in 1 mL cold DMEM/F-12 with Glucose, L-Glutamine and HEPES (Lonza 12-719F). 200 µL of medium containing cells/organoids from the suspension was used for transplantation per mouse. Tumors were transplanted in mice as previously described in accordance with SOP BCCRC 009^36^. Female NOD/SCID/IL2Rγ −/− (NSG) and NOD/Rag1−/−Il2Rγ −/− (NRG) mice were bred and housed at the Animal Resource Centre at the British Columbia (BC) Cancer Research Centre. Disaggregated cells and organoids were resuspended in 150 - 200 µl of a 1:1 v/v mixture of cold DMEM/F12: Matrigel (BD Biosciences, San Jose, CA, USA). 8-12- week-old mice were anesthetized with isoflurane and the suspension was transplanted under the skin on the left flank using a 1 mL syringe and 21-gauge needle. The animal care committee and animal welfare and ethical review committee, the University of British Columbia (UBC), approved all experimental procedures.

### Anchoring, gelation, homogenization and expansion

Fixed cells and tissue slices were first pre-incubated with 100 mM sodium bicarbonate (pH = 8.5, DNase/RNase-free) twice for 15 min each, and incubated in GMA in 100 mM sodium bicarbonate for 3 h at room temperature or 37°C, dependent on target and sample types (detailed anchoring conditions are provided in **Supplementary Table 4**). Of particular note, the solubility of GMA is about 3% in most aqueous solutions and so the anchoring buffer has to be vigorously vortexed after addition of GMA. According to the safety data sheet of GMA, handling of undiluted GMA needs to be done in a fume hood with sufficient ventilation. For experiments using cultured cells, 0.04% (w/v) GMA was used for anchoring, while 0.1% (w/v) GMA was used for tissue samples. After the anchoring reaction, samples were washed with sterile PBS or DPBS three times (for samples using >0.2% GMA in the concentration optimization experiments of **Supplementary Figure 5**, they were washed with 70% ethanol to remove unreacted GMA before washing with DPBS). Then, standard ExM steps including gelation, digestion and expansion were conducted.

Briefly, for gelation, the monomer solution – StockX – was prepared as developed in published protocols^6^: 8.6% (w/v) sodium acrylate (SA), 2.5% (w/v) acrylamide (AA), 0.15% (w/v) N,N’-methylenebisacrylamide (Bis), 2 M sodium chloride (NaCl), 1× PBS. Then the gelation solution was prepared by mixing StockX with 0.5% (w/v) 4-Hydroxy-TEMPO (4-HT) stock solution (required for tissue samples), 10% (w/v) N,N,N′,N′- Tetramethylethylenediamine (TEMED) stock solution, and 10% (w/v) ammonium persulfate (APS) stock solution at 47:1:1:1 ratio on a 4°C cold block, and diffused into the sample at 4°C for 30 min. #0 coverslips were used as spacers between two slides to make a gelling chamber, to cast the gel with thickness around 100 µm. Next, the gelling chamber containing the tissue with infiltrated gelation solution was transferred to a sealed Tupperware for free-radical initiated polymerization at 37°C (a detailed ExM manual can be found here^38^). For the modified 7× expansion protocol, the following monomer solution was used (adapted from the TREx protocol^24^): 17.5% SA, 5% AA, 0.015% Bis, 2 M NaCl, 1× PBS, and mixed with 10% TEMED and APS at 198:1:1 ratio. To reduce the gel attachment to glass surfaces, the glassware can be briefly rinsed with Sigmacote reagent before use.

After 2 hours, the gelled sample was removed from the chamber, trimmed to the proper size, where only the areas of interest were kept, and then immersed in digestion buffer. Different sample homogenization methods were applied as specified below.

For experiments in **Figures 1, 2, 3, Supplementary Figures 4, 5, 6, 10, 11A-B**, the standard proteinase K (proK) based digestion method was performed with the buffer containing 8 U/ml proK, 0.5% (w/v) Triton X-100, 1 mM EDTA, 50 mM Tris-HCl buffer (pH = 8), and 2 M NaCl. The gelled samples were digested at 37°C overnight.

For the comparison experiments of LabelX/AcX and epoxides in preservation of proteins under high heat treatments (**Supplementary Figure 8**), two homogenization methods were tested. In the heat-based SDS denaturation, gels were incubated with the denaturation buffer containing 200 mM SDS, 200 mM NaCl, and 50 mM Tris base (pH = 9) at 95°C for 1 hour, followed by incubation at 37°C overnight and washing with 0.2% TritonX-100 in fresh PBS to remove residual SDS before expansion. In the proK based rapid digestion, gels were digested with 8 U/ml proK, 0.8 M guanidine hydrochloride, 0.5% (w/v) Triton X-100, 1 mM EDTA, 50 mM Tris-HCl buffer (pH = 8), and 2 M NaCl at 60°C for 2 hours, followed by incubation with fresh digestion buffer at 37°C overnight before expansion.

For post-expansion antibody staining and experiments involving WGA staining (**Supplementary Figures 3, 11C, 12**), gelled samples were digested with 50 µg/mL (for cells) or 100 µg/mL (for tissues) endoproteinase LysC in 1 mM EDTA, 50 mM Tris-HCl (pH = 8) and 0.1 M NaCl at 37°C overnight (for cells) or 2-3 days (for tissues). For post-expansion antibody staining targeting Thy1-YFP in mouse brain tissues (**Supplementary Figure 3**), the heat-based SDS denaturation was applied.

After homogenization, the gelled samples were rinsed 3 times with fresh PBS, followed by expansion with ion-free, ultrapure water (3 × 15 min for cells, 3 × 30 min for tissues). To expand LysC-digested tissues, serial incubation with decreasing PBS (1×, 0.5×, 0.1×, 30 min each) or NaCl solutions (1M, 0.5M, 0.1M, 30 min each) was conducted before expansion with water.

### Pre-expansion antibody staining

To assess the potential sample distortion during the GMA-based uniExM procedure and the improvement on imaging resolution, pre-expansion antibody staining was performed. Primary antibodies against β-tubulin, MAP2, neurofilament, GFP, and βII- spectrin were used to stain predetermined structures in different samples. In brief, samples were fixed with 3% PFA/0.1% GA (for microtubule and spectrin preservation) or 4% PFA, followed by processing with 0.1% sodium borohydride and 100 mM glycine to quench unreacted fixative residuals. MAXblock medium was used for blocking for 1 hour and then 5 μg/mL primary antibody diluted in MAXbinding medium was incubated with the sample at 4°C overnight (or 37°C for 2 hours). Next day, 5 μg/mL fluorescently labelled secondary antibody was used at room temperature for 1 hour. After completely washing out unbound antibodies, samples were proceeded with the anchoring and expansion steps.

### Post-expansion antibody staining

For post-expansion antibody staining, gelled specimens were digested with the milder enzyme LysC and expanded^3^. Then the samples were incubated with 5 μg/mL primary antibody diluted in MAXbind staining medium at 4°C overnight (or 37°C for 2 hours) and washed with MAXwash medium four times. 5 μg/mL fluorescently labelled secondary antibody was incubated with the sample to develop signals before DAPI counterstaining and expansion.

### Staining for lipids and carbohydrates

R18, FM and BODIPY were all tested for pre- and post-expansion staining. The working concentration for these dyes was chosen to be 10 μg/mL (diluted with fresh PBS). For pre-expansion staining, lipid tags were introduced right before the gelation step in which samples were stained for 1-2 hours at room temperature. Then the samples were anchored with GMA at 4°C overnight, followed by incubation at room temperature for another 3 hours, digestion (proK-based) and expansion. For post-expansion staining, samples were fixed with 3% PFA/0.1% GA, and then anchored with GMA. The samples were digested with proK or LysC (for WGA staining), followed by expansion. After expansion, the samples were stained with antibodies or HCR-FISH first (if at all), and then with lipid tags or WGA-A647 for 1-2 hours at room temperature. The working concentration for WGA-A647 was chosen to be 5 μg/mL. Residual dyes were washed off with 1% Zwittergent in DPBS.

### Expansion fluorescence *in situ* hybridization (ExFISH)

For ExFISH experiments (in **Figure 1B, Supplementary Figures 3, 6, 10B**) expanded gels were re-embedded in 3% AA-based (plus 0.15% BIS, 5 mM Tris base pH 9, and 0.05% APS/TEMED), non-expandable gel to maintain rigidity. With the re-embedding step, the expansion factor would decrease to ∼3.2 compared to the original expansion factor of ∼4.2. Two stacked #1.5 coverslips were usually used as the spacers between two glass slides for re-embedding. The HCR-FISH probes and reagents were purchased from Molecular Instruments, Inc. In general, the gel was incubated with hybridization buffer at room temperature for 30 min, and then with 1:500 diluted gene-specific probe (8 nM total final probe concentration) set at 37°C overnight. Next day, the gel was washed with HCR washing buffer at 37°C for 4 × 30 min and with 5× SSCT buffer (5× SSC buffer containing 0.1% Tween 20) at RT for 4× 15 min, followed by incubation with 1:200 diluted, fluorescently labelled HCR hairpin amplifiers at room temperature overnight. Lastly, the gel was washed with 5× SSCT for 4 × 20 min and counterstained with 1 μg/mL DAPI. To characterize RNA capture efficiency by GMA, HCR-FISH against the same genes was performed in the same sample before and after anchoring. Before anchoring, HCR-FISH was done in HeLa cells and then the hybridized probes were removed with 80% formamide. Then, ExFISH after GMA-based uniExM was done with the same sample, where the same cells were imaged in both conditions. Transcripts in single cells were counted using MATLAB scripts as developed before^39, 40^.

### Expansion Sequencing (ExSeq)

The detailed protocol for ExSeq was published previously and involves a multi-day procedure^21^. 87 target genes were chosen to differentiate two major cancer clones in the SA501 PDX line (full gene list is provided in **Supplementary Table 5**)^37^. In brief, a re-embedded gel was passivated with 2 M ethanolamine, 150 mM EDC and NHS. Then the passivated gel was subjected to targeted ExSeq (tExSeq) or untargeted ExSeq (uExSeq). For tExSeq, padlock probes targeting specific mRNAs (in general, 12-16 probes per gene and 100 nM per padlock probe diluted in 2× SSC containing 20% formamide) were used to hybridize with the sample at 37°C overnight. Then the unhybridized probes were completely washed off with fresh hybridization buffer and the sample was treated with 1.25 U/μL PBCV-1 DNA ligase at 37°C overnight, followed by inactivation at 60°C for 20 min. Next, the successfully ligated padlock probes were rolling circle amplified with 1 U/μL phi29 DNA polymerase at 30°C overnight. As all padlock probes targeting the same gene bear a predetermined barcode, the identity of the mRNA can be read out by commercially available sequencing reagents (*e.g.,* the Illumina MiSeq kit). In comparison, uExSeq utilizes randomized 8N oligonucleotide probes to hybridize with any potential RNA targets without prior sequence knowledge.

After that, reverse transcription was performed *in situ* with 10 U/μL SSIV reverse transcriptase to generate cDNAs containing inosine. The cDNAs were later segmented to proper sizes with endonuclease V and circularized with 3 U/μL CircLigase. Then the target mRNAs were digested away with RNase H. Such circularized cDNAs were subjected to rolling circle amplification and sequencing readout. For detailed working mechanism and protocols of tExSeq and uExSeq, please refer to our previous work^21^.

We adapted the sequencing-by-synthesis chemistry for *in situ* 7-base readout using the Illumina MiSeq v3 kit with a modified protocol. To help the registration process, the re-embedded gel sample was adherent to bind-silane (1:250 diluted in 80% ethanol) processed glass surface with the same re-embedding monomer solution containing 1:100 diluted TetraSpeck microspheres. Before sequencing, the sample was first treated with 400 U/mL terminal transferase and 50 µM ddNTP to block nonspecifically exposed 3’ ends in DNA, and then hybridized with 2.5 µM sequencing primer (5’-TCT CGG GAA CGC TGA AGA CGG C-3’) in 4× SSC at 37°C for 1 hour. After 3 × 10 min washing with fresh 4× SSC, the sample was incubated with the PR2 incorporation buffer (part of the MiSeq kit) for 2 × 15 min. Then the sample was pre-incubated with 0.5× incorporation mix buffer (IMT of the MiSeq kit) supplemented with 1× Taq polymerase buffer and 2.5 mM magnesium chloride at RT for 2 × 15 min. Then the sample was incubated with 0.5× IMT at 50°C for 10 min for one base elongation. After the elongation reaction, the sample was washed with PR2 containing 2% Zwittergent at 50°C for 2 × 15 min followed by additional washing with PR2 at RT for 2 × 15 min. Next, the sample was immersed in imaging buffer (SRE of the MiSeq kit) and subjected to imaging (elaborated in the following section). After imaging, the sample was briefly washed with PR2 at RT for 2 × 10 min. Then the sample was incubated with cleavage solution (EMS of the MiSeq kit) at 37°C for 3 × 15 min. Lastly, the sample was washed with PR2 at 37°C for 2 × 15 min and at RT for 2 × 15 min, and then started with the next round of elongation process.

### Data analysis for ExSeq

Data analysis for the sequenced PDX sample followed our established ExSeq processing pipeline (available at: https://github.com/dgoodwin208/ExSeqProcessing). For the 87-gene probe set, a 7-base barcoding strategy with error correction capacity was adopted. Upon microscopic readout, the raw image files were stored in 16-bit HDF5 format and subjected to color correction, registration, segmentation, basecalling and alignment as done in our previous work,^21^ and then performed manual cell segmentation in 2D according to a max-Z projection of the DAPI staining channel using the VASTLite package (https://lichtman.rc.fas.harvard.edu/vast/). In total, 793,535 unique transcripts were detected for a population of 3,339 cells (with effective lateral resolution ∼78 nm and axial resolution of ∼160 nm). For gene function annotation we refer to The Human Protein Atlas (https://www.proteinatlas.org) or The Human Gene Database (https://www.genecards.org). The spatial maps of single transcripts or functional gene groups were generated with MATLAB scripts (for coordinates extraction) and ImageJ packages (for visualization).

For single-cell biostatistics analysis we utilized the R toolkit Seurat 4. Before analysis, we further pruned the dataset based on the counts per cell values, where cells with less than 50 counts or more than 3000 counts were filtered out and we ended up with 2,732 cells. Then we normalized the counts per cell by the median value from all the cells and performed a log transformation. To identify cell clusters, we applied both unsupervised and supervised approaches. In unsupervised clustering, PCA suggested a majority of cells could be classified into two groups using the expression of 30 genes automatically pulled out by PCA. With that, we decided to visualize these two cell groups based on the relative expression of these 30 genes (by correlating the percentages of the top and bottom 15 genes in each single cell with the color channel intensities of an RGB composite image). Therefore, for every cell, its closeness to a particular group rather than an arbitrary binary classification was presented so that the transitional status of different tumor clones may be preserved. The gene list used for supervised clustering was selected based on the bulk RNA-seq expression data, in which a set of 15 marker genes for each clone (Clone_ZNF24: *RNF146, DDX24, OAZ2, ZNF24, TXNL1, IDH2, SEPT4, CDCA7, CP, RAD21, WDR61, RBP1, COX5A, HSPE1, IER3IP1*; Clone_XIST: *XIST, CD44, FBXO32, LGALS1, ARC, HLA-A, HLA-C, S100A11, CTSV, SLC25A6, ANXA1, ARHGDIB, SQLE, B2M, NDUFS5*) in the SA501 PDX model was applied for the initial dimension reduction. Afterwards, the major cell clusters were presented via uniform manifold approximation and projection (UMAP). To cross-check the agreement between unsupervised and supervised cell classification results, we randomly sampled 2,000 cells from the two tumor clones as annotated by either cell classification method, and determined if they were correctly assigned to the same clone by the other cell classification method (in unsupervised cell classification, a minimum level of 30% for the summed 15 marker gene counts to total gene counts was applied as the threshold for robust clone assignment).

### Imaging and image analysis

All imaging experiments were performed on a spinning disk confocal microscope (Andor Dragonfly) equipped with a Zyla sCMOS 4.2 plus camera (pixel size 6.5 μm) or a CSU- W1 SoRa super-resolution spinning disk confocal microscope (Nikon). Six main lasers on Dragonfly were used: 405 nm (100 mW), 488 nm (150 mW), 561 nm (150 mW), 594 nm (100 mW), 637 nm (140 mW) and 685 nm (40 mW). For tiled scan of full-size brain slices (**Figure 1B, i; Supplementary Figure 8**), a 10× objective lens was used. For other imaging experiments, a Nikon CFI Apochromat LWD Lambda S 40XC water immersion objective lens (working distance 0.6 mm, NA 1.15) was used together with the Zeiss Immersol medium (Refractive Index 1.3339). For ExSeq using Illumina Miseq reagents, the following bandpass filters were used: 705-845 nm for base “C” channel, 663-737 nm for base “A” channel, 575-590 nm for base “T” channel, 500-550 nm for base “G” channel. All channels used 200 ms as the exposure time except that base “G” used 400 ms exposure time. For the sequenced cancer tissue, a total of 12 × 6 fields of view (FOV dimension in pre-expansion units: 104 × 104 × 62.5 µm^3^) were captured.

For characterization of expansion in uniExM, HeLa cells stained with β-tubulin antibody and DAPI were used. The size of cells was represented by the largest diameter value from the microtubule staining image, and this measurement was performed on the same cells pre- and post-expansion. In parallel, the area and shape descriptors of cell nuclei were measured with ImageJ. The following four parameters were obtained:

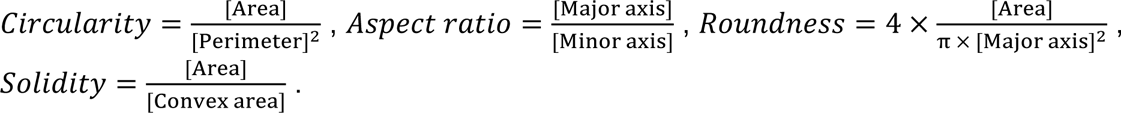

Quantification of expansion errors was performed as previously described^2, 3^. In brief, HeLa cells were stained with β-tubulin antibody pre-expansion. The same cells were imaged both pre- and post-expansion, where the pre-expansion images were taken with a Nikon SoRa super-resolution microscope (∼1.8-time spatial resolution improvement over standard confocal microscopy). The obtained images were first histogram normalized and deconvolved in imageJ. Then non-rigid registration was performed using B-spline grids to capture potential non-uniformities between images.

For periodicity analysis of βII-spectrin, cultured neurons were stained pre-expansion. Then the cells were expanded and imaged. Segments of axon processes with more than 10 spectrin signal clusters (a signal cluster was defined as fluorescence intensity above 5 times of the standard deviation of the background level) were selected and relevant fluorescence profiles were extracted. The fluorescence traces in space were scaled back to the pre-expansion level and autocorrelation was performed in OriginLab software. From the obtained autocorrelation curve, periodicity was calculated by averaging the distances of the first four adjacent peaks.

## Acknowledgements

We acknowledge Asmamaw T. Wassie, Tay Shin, Ruihan Zhang and other Boyden lab members for insightful discussions. E.S.B. acknowledges for funding, Lisa Yang, John Doerr, Open Philanthropy, Tom Stocky, NIH 1R01EB024261, Kathleen Octavio, Lore McGovern, NIH 1R01AG070831, HHMI, NIH 1R01MH123403, ERC Synergy Grant agreement No 835102, NIH R01MH124606, NIH R37MH080046, NIH U24NS109113, NIH 1U19MH114821, and Cancer Research UK Grand Challenge grant C31893/A25050.

## Author contributions

Y.C., G.Y. and E.S.B. initiated the study and contributed to the key ideas. Y.C. and G.Y. screened candidate anchor molecules and performed proof-of-concept experiments for uniExM. Y.C. designed and performed experiments, analyzed and interpret the data for uniExM in RNA quantification, protein ultrastructure analysis and multimodal detection.

Y.C. and G.Y. co-developed the protocols for uniExM in lipids and carbohydrates detection. Y.C., D.R.G., C.H.O., A.S. and S.A. designed and performed experiments, analyzed and interpreted the data for uniExM in *in situ* sequencing and PDX breast cancer tissue mapping. C.Z. and K.E.K. contributed to the conceptualization of multimodal ExM. Y.C. and C.Z validated and optimized the expansion protocol. K.E.K. and D.P. helped animal surgery and brain sample preparation. Y.C. and E.S.B. wrote and edited the manuscript, with critical input from all authors. E.S.B. supervised the study.

## Competing interests

Y.C., G.Y. and E.S.B. have filed for provisional patent protection on the design, steps and applications of uniExM through MIT Technology Licensing Office (U.S. Appl. No. 63/326,346). E.S.B. is a cofounder of the company Expansion Technologies to help with commercialization of ExM related techniques.

## Supplementary Figures

**Supplementary Figure 1.**
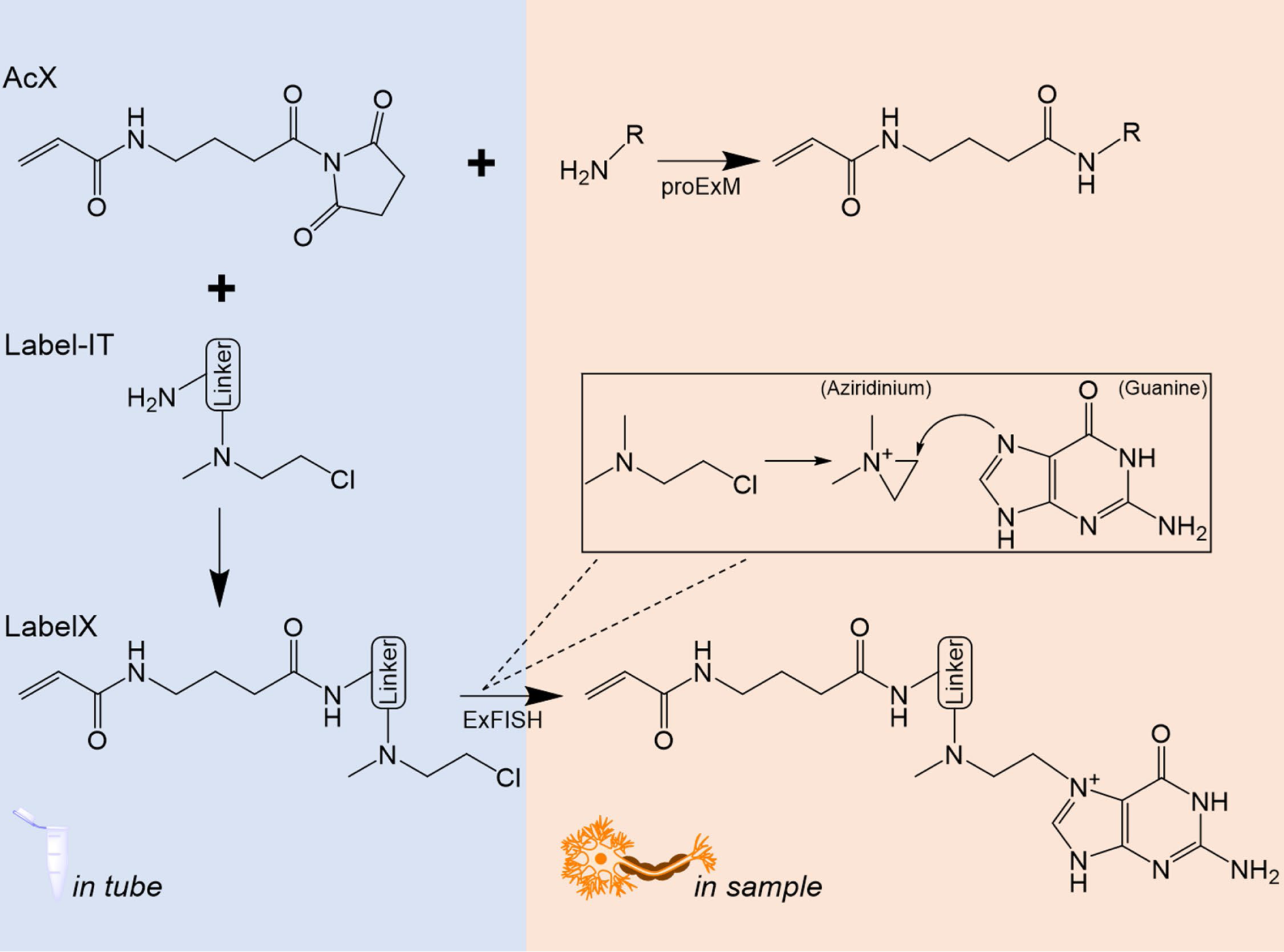
Working mechanisms of the anchoring chemistry of standard ExM. In protein-retention ExM (proExM), AcX functions by modifying the amine groups on proteins through a succinimidyl ester moiety. The acryloyl group of AcX crosslinks the proteins to the polyacrylate hydrogel. In order to anchor nucleic acids, AcX is first reacted with an alkylating reagent such as Label-IT amine to form aziridinium-containing LabelX which can later be coupled to the N7 position of guanine in DNA and RNA. However, this in-house synthesis of anchor molecules suffers from high cost, nonstandard yield, and increased procedure time and complexity.

**Supplementary Figure 2.**
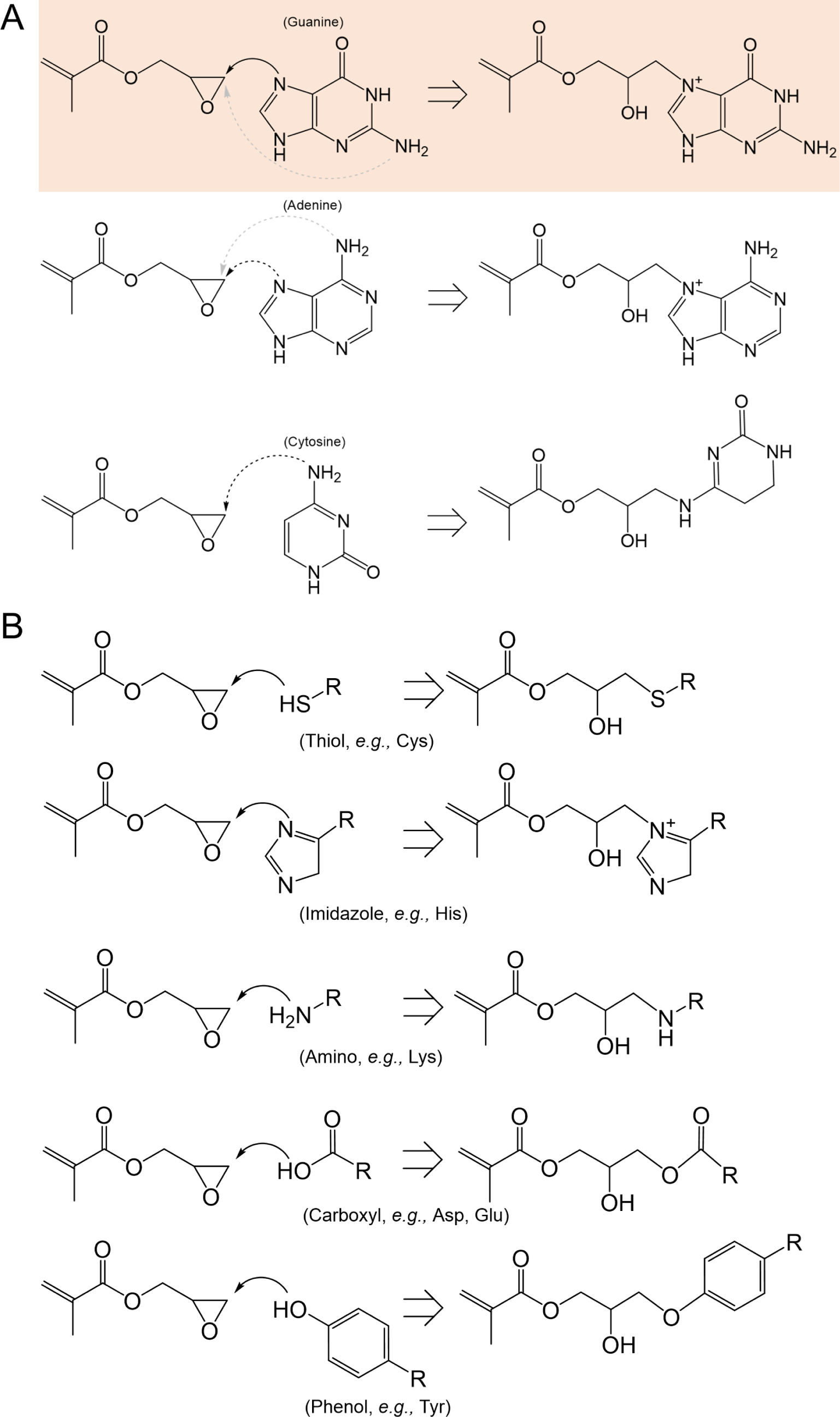
Representative potential nucleophile substrates in a biological system, including **(A)** nucleic acids and **(B)** amino acids. A potential reaction between GMA and N7-guanine in anchoring DNA and RNA is highlighted in orange. Potential reactions between common nucleophilic amino acids and GMA are illustrated.

**Supplementary Figure 3.**
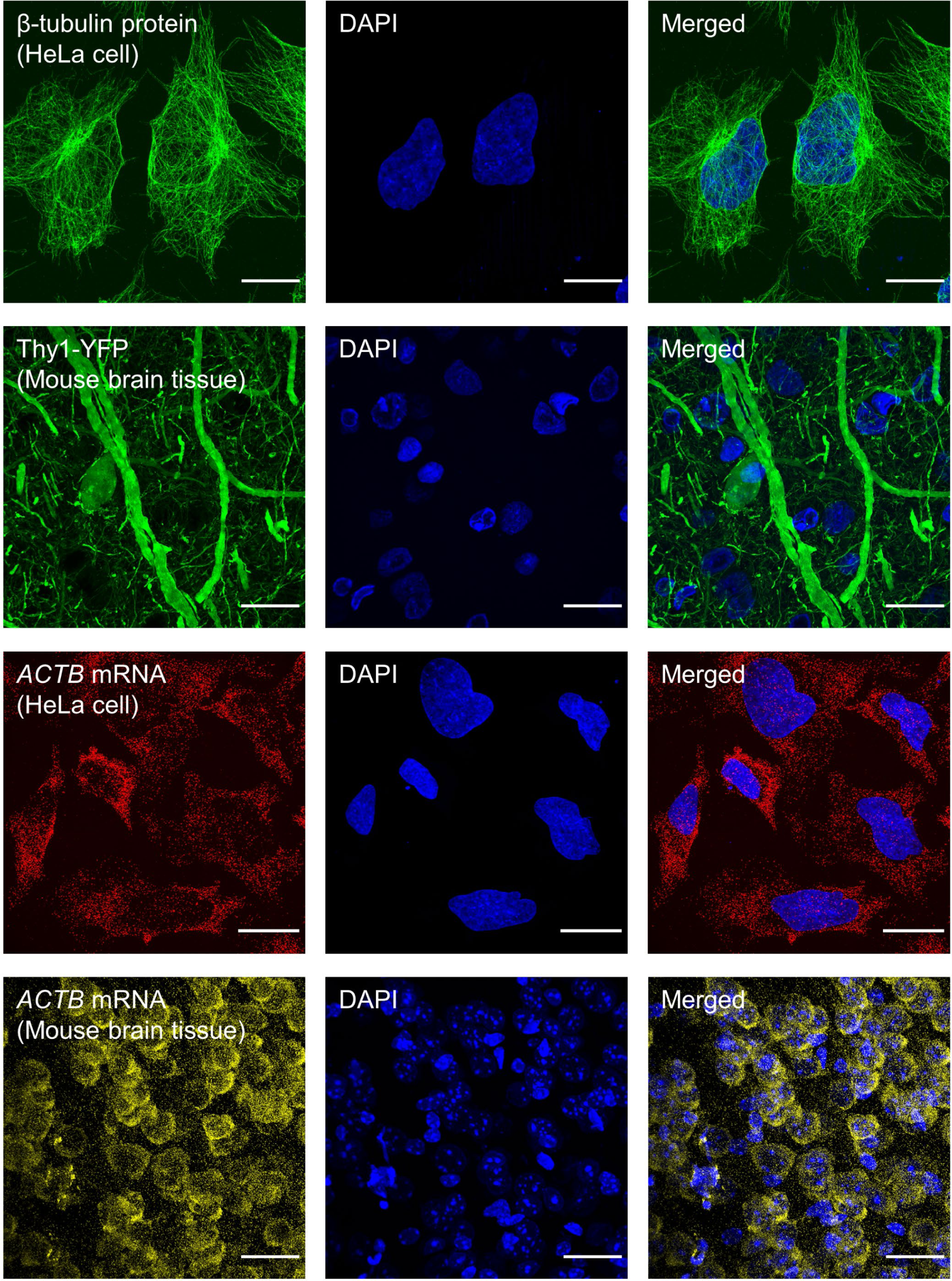
GMA-based uniExM enables retention of proteins and RNAs in expanded cells and tissues. In protein retention tests, β-tubulin in HeLa cells and YFP in mouse brain tissues were chosen as targets. Row 1: HeLa cells were anchored with 0.04% (w/v) GMA, digested with LysC and stained with anti-β-tubulin (and DAPI) post-expansion. Row 2: A 50 µm thick Thy1-YFP mouse brain slice was anchored with 0.1% (w/v) GMA, digested with proK, and imaged post-expansion. In RNA retention tests, HCR-FISH targeting *ACTB* in HeLa cells (Row 3) and mouse brain tissues (Row 4) were performed post-expansion, respectively. Color representation in the images: blue – DAPI; green – Alexa488; yellow – Alexa546; red – Alexa647. Scale bars (in pre-expansion units): 20 µm.

**Supplementary Figure 4.**
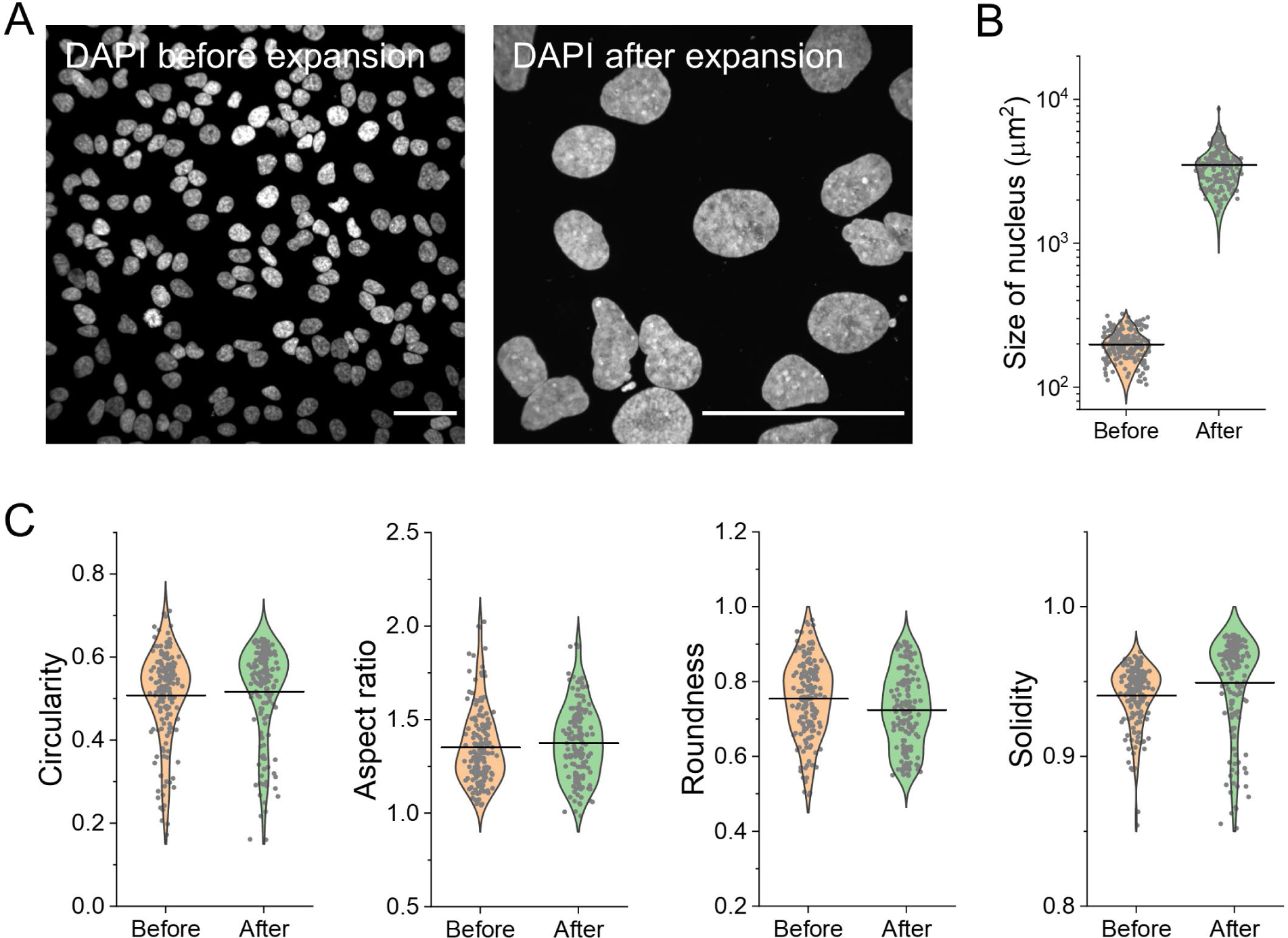
Assessment of the size and morphological properties of HeLa cell nuclei before and after GMA-based uniExM. **(A)** Representative images of DAPI staining for HeLa cell nuclei before and after expansion. Scale bars (in pre-expansion units): 50 µm. **(B)** Size of nuclei was measured before and after expansion. **(C)** 4 parameters related to nuclear morphological properties – circularity, aspect ratio, roundness and solidity – were evaluated within ImageJ. (Data presented in violin plots with raw data points shown and mean values highlighted with solid lines, n = 200 cells from 4 culture batches, two sample *t*-test was performed with all *p* > 0.1)

**Supplementary Figure 5.**
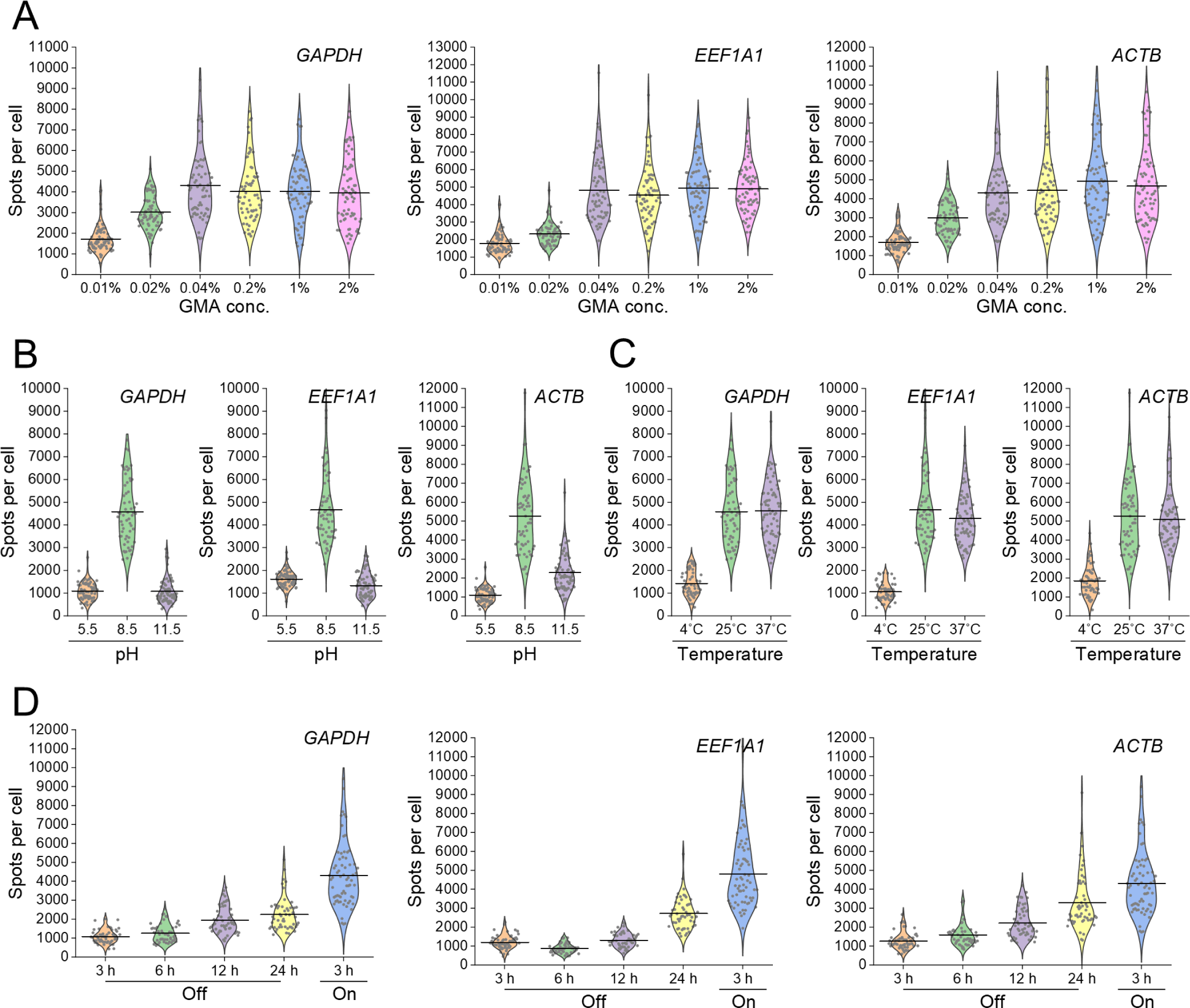
Optimization for GMA-based RNA anchoring in uniExM. **(A)** Different concentrations of GMA, **(B)** anchoring pH and **(C)** temperatures were tested in the context of ExFISH targeting three genes – *GAPDH, EEF1A1 and ACTB* – in HeLa cells, to determine optimal RNA retention conditions. **(D)** In light of the above tests, the GMA anchoring reaction for RNAs could be tuned “On” and “Off” by varying the reaction temperature and pH. Reaction “Off” condition: 4°C, pH = 7.4 with 1×PBS. Reaction “On” condition: 25°C, pH = 8.5 with 100 mM NaHCO3. To test the “Off-On” transition, one group of cells were first incubated under the “Off” condition for 12 h, followed by treatment under the “On” condition for 3 h. (n = 50-80 cells for each tested condition and from 2 culture batches)

**Supplementary Figure 6.**
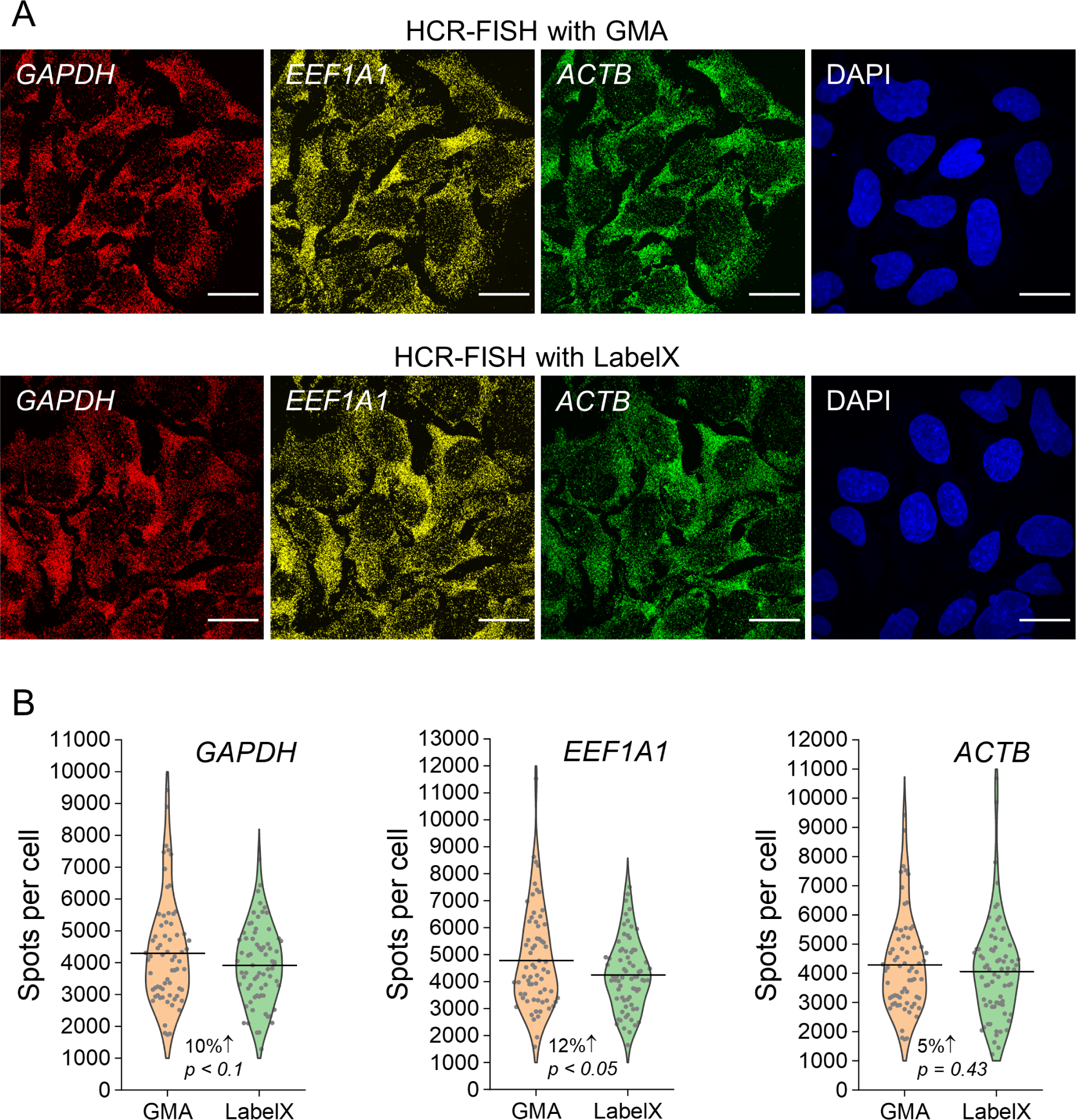
Head-to-head comparison between GMA and LabelX in RNA retention and detection. **(A)** HCR-FISH targeting three highly expressed housekeeping genes was performed in HeLa cells treated with 0.04% (w/v) GMA or 0.01% (w/v) LabelX, under their respective optimal conditions. Color representation in the images: blue – DAPI; green – Alexa488; yellow – Alexa546; red – Alexa647. Linear expansion factor: 4.3 for GMA; 4.4 for LabelX. Scale bars (in pre-expansion units): 20 µm. **(B)** Summary plots of detected transcripts per cell for each gene. (Data presented in violin plots with raw data points shown and mean values highlighted with solid lines; n = 70-100 cells from 3 culture batches; two-sample *t-*test was performed, with *p* values shown)

**Supplementary Figure 7.**
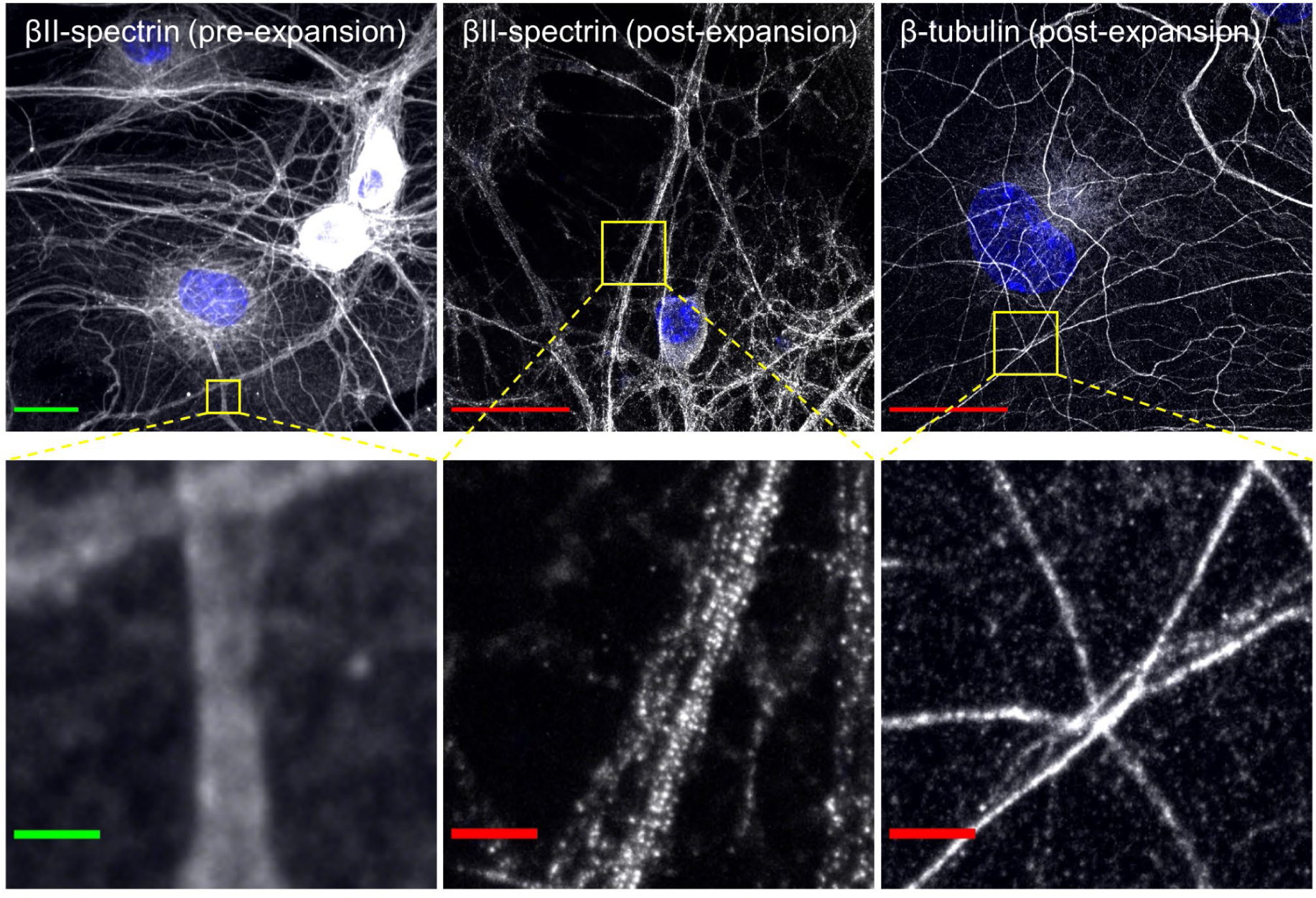
uniExM helps resolve the ultrastructure of βII-spectrin in neuron axons – additional images. Left column: antibody staining for βII-spectrin in mouse hippocampal neurons shows apparently continuous signal distribution along neuronal processes. Middle column: the periodic, punctate distribution of βII-spectrin signals is revealed, after expansion. Right column: antibody staining against β-tubulin shows continuous microtubule structures even with expansion. All antibody staining was performed pre-expansion. Linear expansion factor: 4.4. Color representation in the images: gray/white – antibody staining; blue – DAPI. Scale bars (in pre-expansion units): 20 µm (upper panels); 2 µm (lower panels).

**Supplementary Figure 8.**
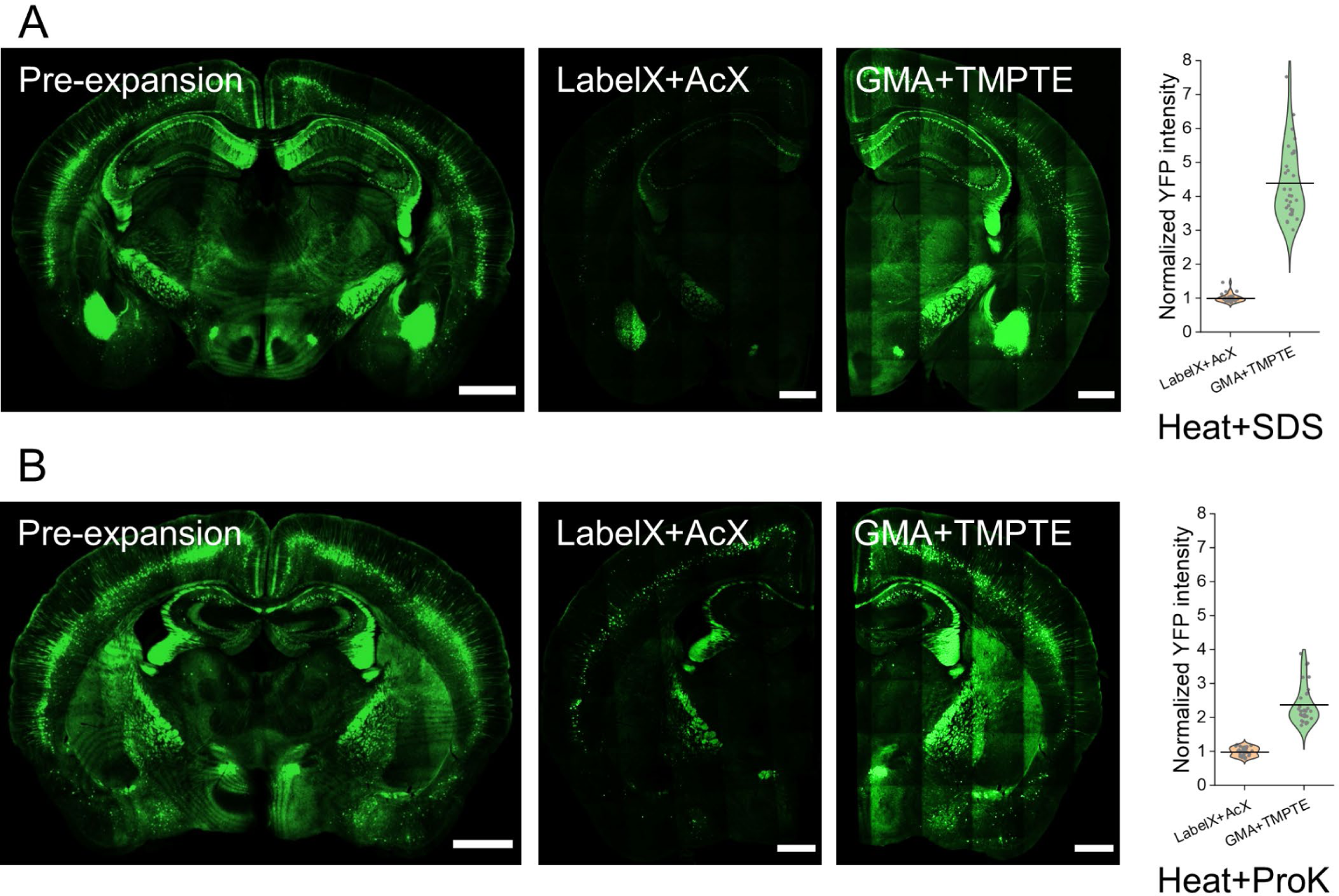
Preservation of fluorescent protein function in tissues undergoing high heat treatments by combining acrylate epoxides and polyepoxides. Retention of Thy1-YFP fluorescence signals in mouse brain tissues by LabelX plus AcX vs. GMA plus TMPTE after: (A) strong detergent SDS-based denaturation: 95°C for 1 h, followed by 37°C overnight; (B) ProK-based digestion: 60°C for 2 h, followed by 37°C overnight. Scale bars (in pre-expansion units): 1,000 µm (whole brain); 500 µm (half brain). Raw intensity measurements are presented in violin plots with mean values highlighted with solid lines. The intensity values were normalized to the level of LabelX plus AcX treated samples. (n = 30 measurements from 4 brain slices, 2 mouse brains; two-sample *t*-test was performed for statistical significance tests, with both *p* < 10^-15^)

**Supplementary Figure 9.**
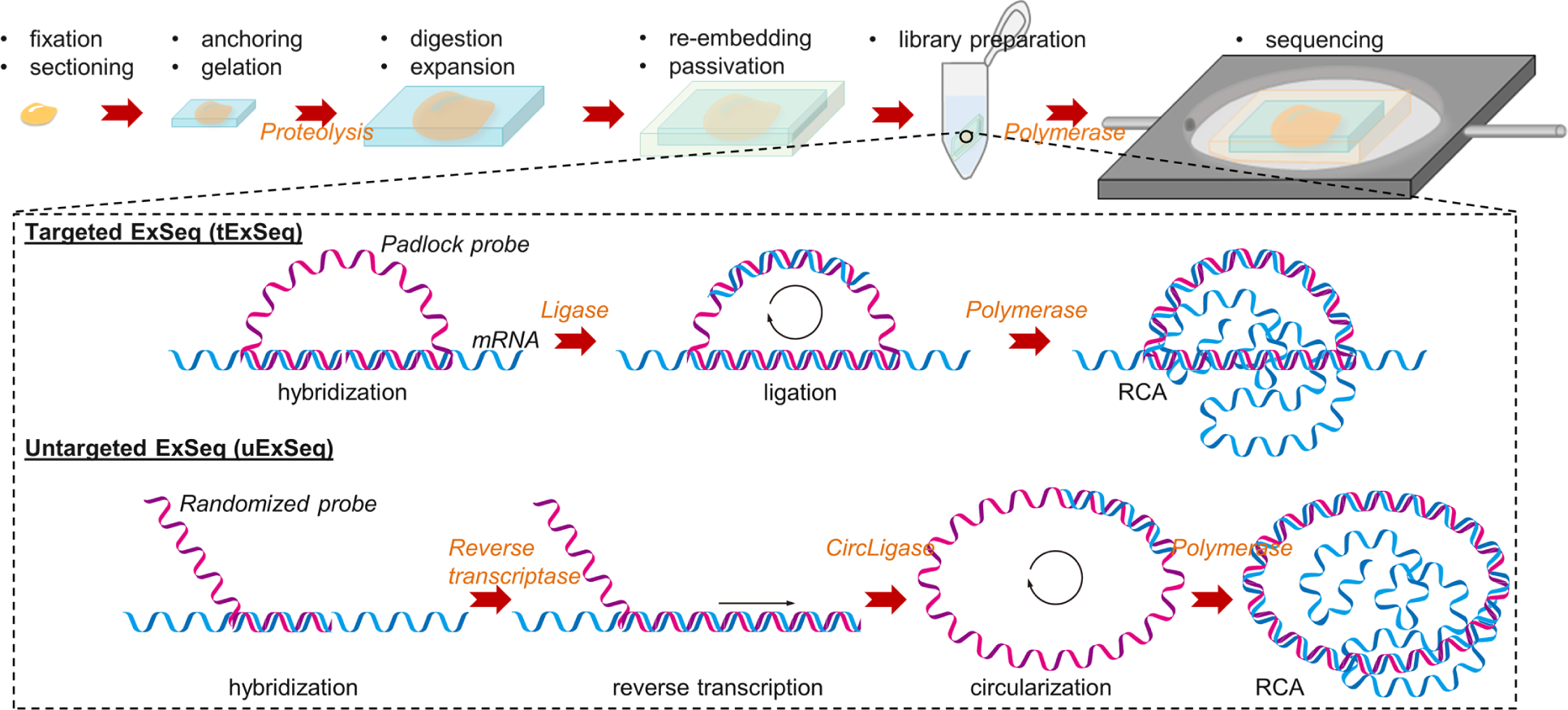
Schematic illustration for targeted and untargeted ExSeq procedures. After a sample is fixed, sectioned, and expanded, it undergoes a second gelation in a charge-neutral hydrogel (the process of “re-embedding”) followed by neutralization of charge on the original hydrogel (the process of “passivation”) to prepare the sample for sequencing. Then padlock probes targeting specific mRNAs (for tExSeq) or randomized oligonucleotide probes (for uExSeq) are introduced. In tExSeq, the padlock probes are directly ligated upon hybridization to their designated targets, while in uExSeq the randomized probes prime reverse transcription to add sequence information from the bound RNA into cDNA form, followed by probe circularization. The ligated or circularized probes are then subjected to rolling circle amplification (RCA) before being sequenced by ligation or synthesis.

**Supplementary Figure 10.**
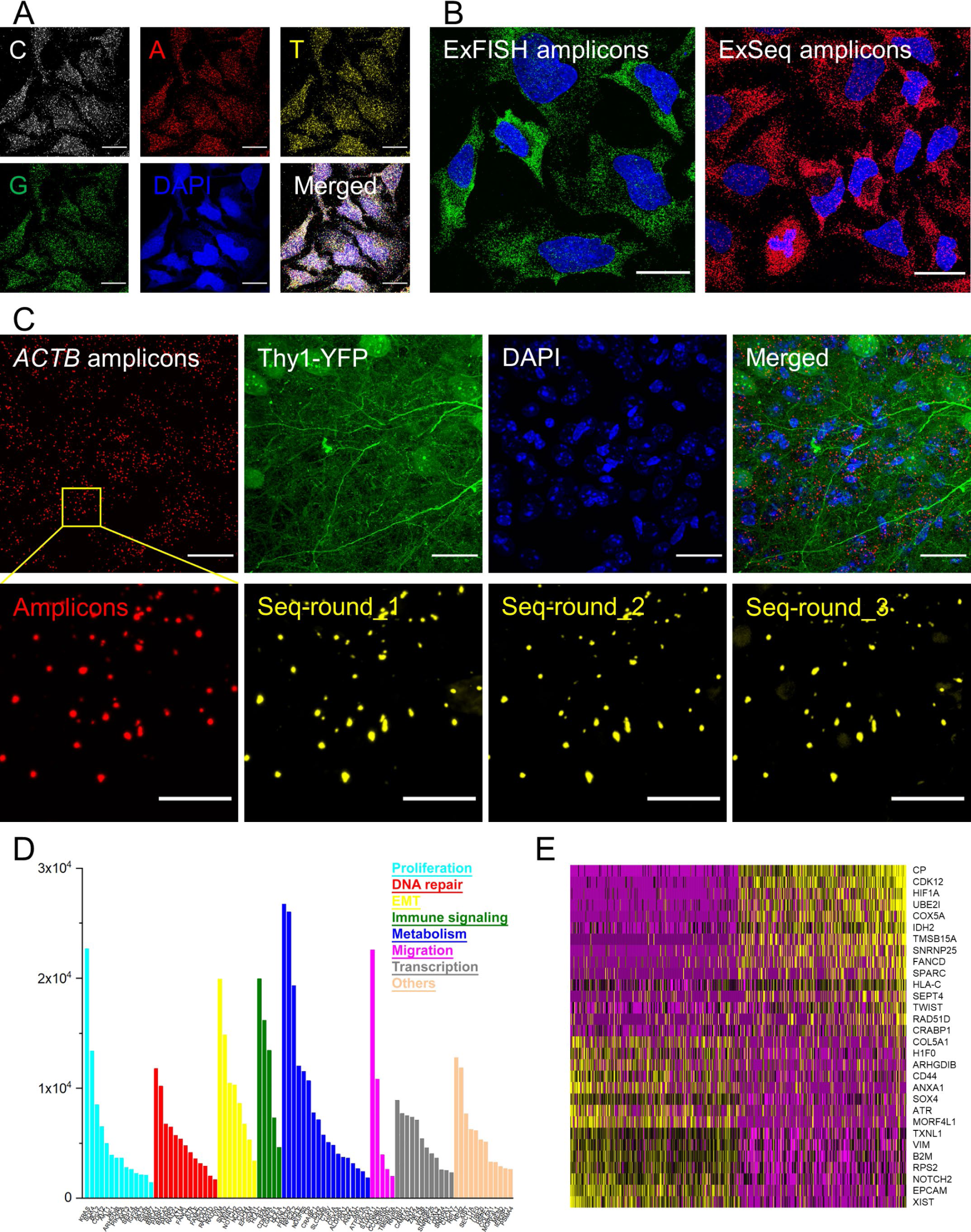
Demonstration of uniExM for *in situ* RNA sequencing (ExSeq). **(A)** Amplicons generated by GMA-based uExSeq in HeLa cells were imaged with SBS reagents (from the Illumina MiSeq v3 kit). The following excitation and emission wavelengths were used for 5-channel acquisition: DAPI – Ex. 405 nm / Em. 440-460 nm; Base “G” – Ex. 488 nm / Em. 500-550 nm; Base “T” – Ex. 561 nm / Em. 575-590 nm; Base “A” – Ex. 640 nm / Em. 663-737 nm; Base “C” – Ex. 685 nm / Em. 705-845 nm. Scale bars (in pre-expansion units): 20 µm. **(B)** Characterization of uniExM for *in situ* enzymatic amplification in tExSeq. *GAPDH* mRNAs were amplified by HCR-FISH (ExFISH) or padlock probes in tExSeq. Then the generated signal spots in individual cells were counted. For better comparison, Alexa546 conjugated oligonucleotide probes were used for amplicon detection in both cases. Scale bars (in pre-expansion units): 20 µm. **(C)** Characterization of uniExM for *in situ* enzymatic sequencing in tExSeq. tExSeq targeting *ACTB* mRNAs in Thy1-YFP mouse brain tissues was performed, where padlock probes bearing consecutive bases “TTT” as the barcode were used. Before *in situ* sequencing, imaging with universal amplicon detection probes was performed to establish a reference image for the transcript locations (lower left, red). YFP signals were also imaged. After that, the universal probes and YFP signals were removed by concentrated formamide and heat treatment. Next, three rounds of SBS were conducted and the detected signal spots were benchmarked against the reference amplicon image (lower row, yellow dots). Scale bars (in pre-expansion units): 20 µm (for upper panel), 5 µm (for lower panel). **(D)** tExSeq targeting 87 cancer clone-specific genes in SA501 PDX breast cancer tissue was performed using 7-round SBS. All the decoded transcripts from a ∼0.8 mm^2^ tissue slice, along with their function annotations, are summarized in the bar chart. **(E)** Principal component analysis (PCA) identified two groups of genes (15 each) that classify the tissue cells into two primary groups.

**Supplementary Figure 11.**
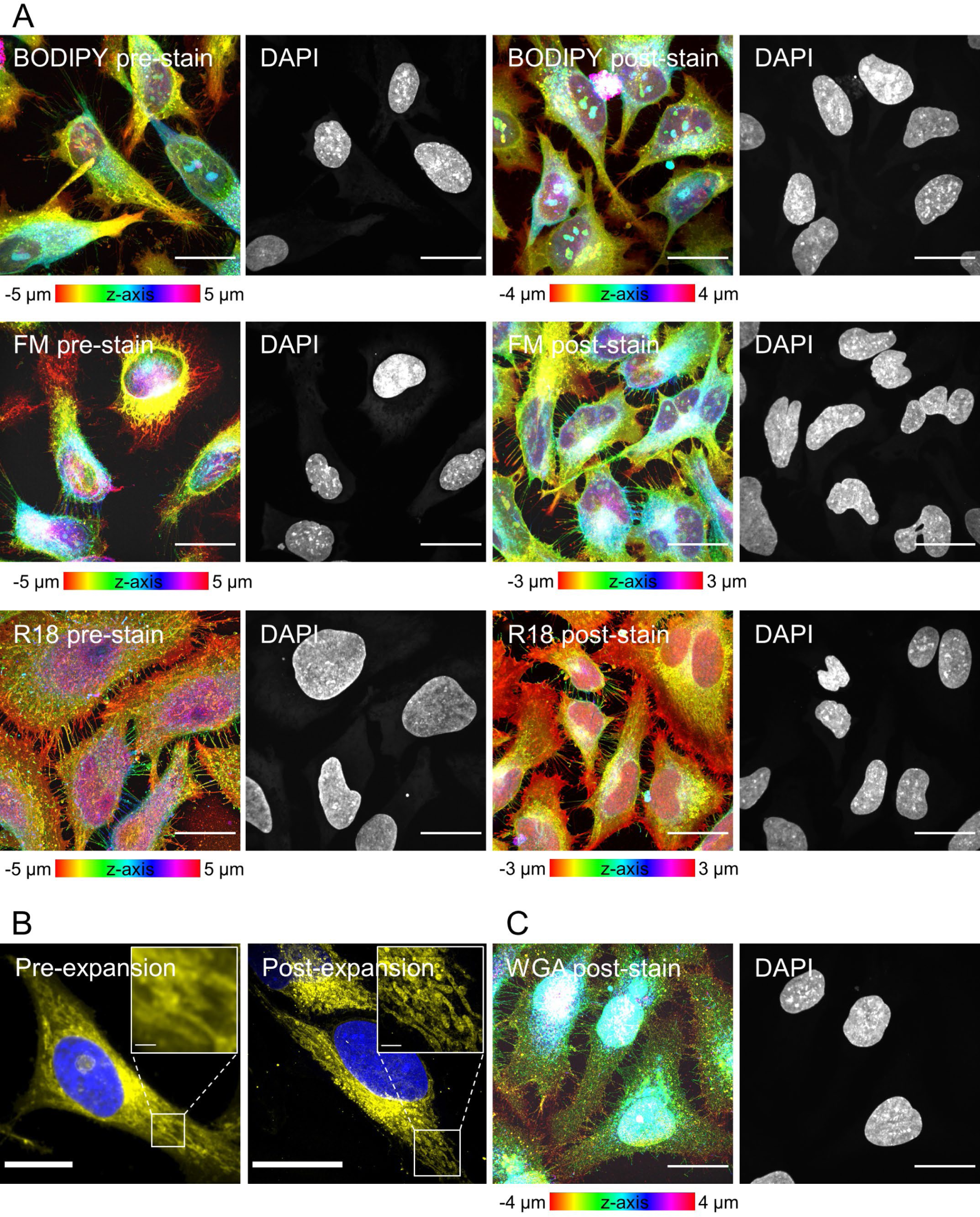
Detection of lipids and carbohydrates in uniExM. **(A)** Commercially available lipid staining reagents – BODIPY, FM and R18 – were tested in HeLa cells in the context of both pre- and post-expansion staining. As shown here, all three staining reagents exhibit consistent image patterns in pre- and post-expansion staining. The color-code in the lipid signal channel represents z-axis information. Scale bars (in pre-expansion units): 20 µm. **(B)** Detection of lipids expands the biological information that can be obtained by uniExM. In the expanded sample, post-expansion staining with lipophilic fluorophore R18 helps better resolve lipid-rich mitochondria where cristae are only discernable with expansion (highlighted in the zoomed-in inset). Scale bars (in pre-expansion units): 20 µm (large images); 1 µm (insets). **(C)** WGA- A647 was used to stain carbohydrates/glycoconjugates post-expansion in HeLa cells digested with the selective protease LysC. Strong signals were detected on cell and nuclear membranes. Colors in the WGA channel represents z-axis information. Scale bars (in pre-expansion units): 20 µm.

**Supplementary Figure 12.**
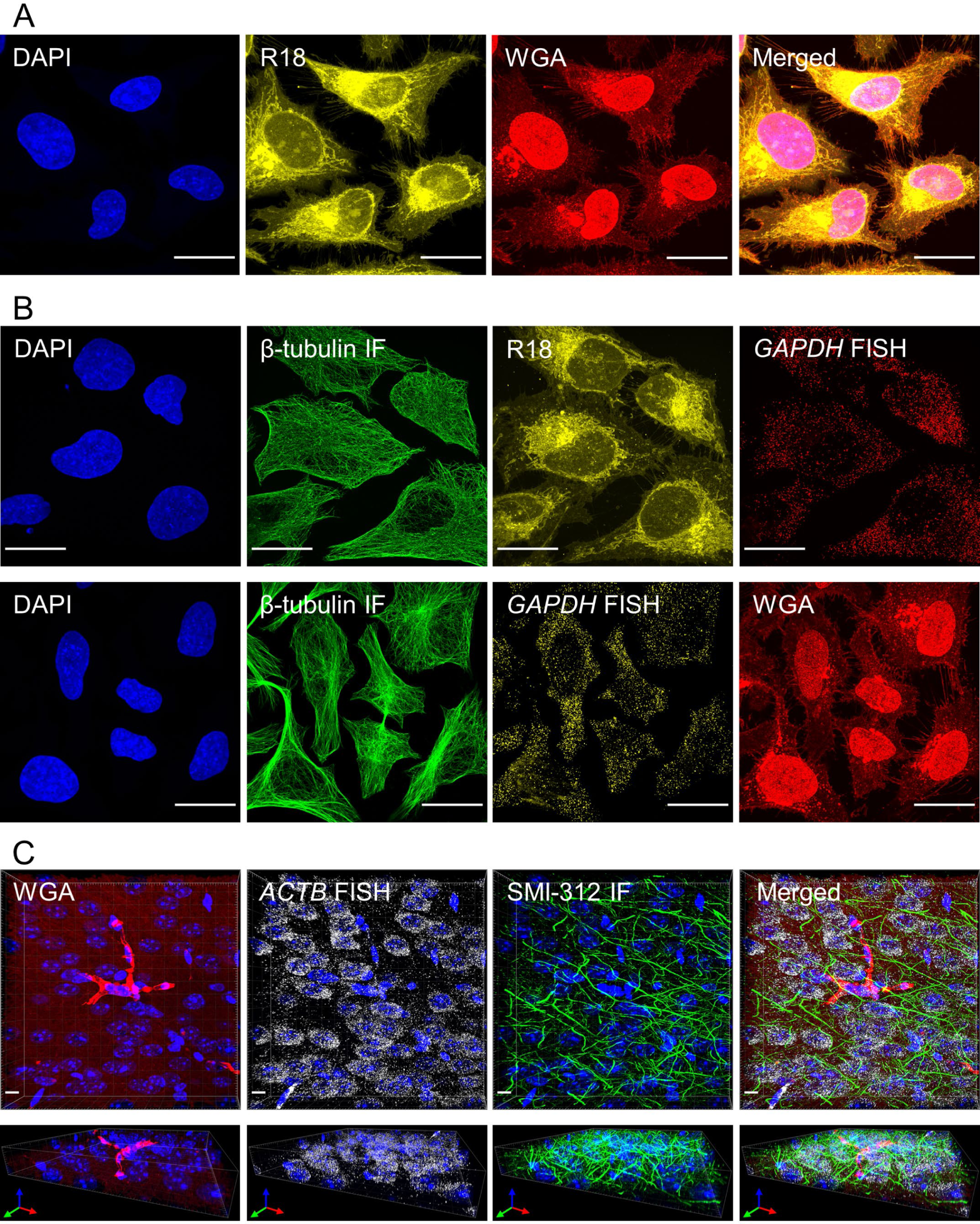
uniExM supports multimodal imaging of multiple biomolecular species. **(A)** Demonstration of post-expansion co-staining for carbohydrates and lipids. HeLa cells were processed with 0.04% (w/v) GMA and LysC proteolysis, followed by staining with 10 µg/mL R18 and 5 µg/mL WGA-A647. Linear expansion factor: 4.3. Scale bars (in pre-expansion units): 20 µm. **(B)** Lipid or carbohydrate staining can be combined with protein and RNA detection in the same sample. As demonstrated in this figure, β-tubulin was stained with antibody pre-expansion, while R18, WGA staining and HCR-FISH were performed post-expansion. In addition to target-specific detection by IF (immunofluorescence) and FISH, staining for lipids and carbohydrates provides structural information at the cellular (*e.g.,* membranes) or subcellular levels (*e.g.,* mitochondria). Scale bars (in pre-expansion units): 20 µm. **(C)** Demonstration of multimodal detection by uniExM at the tissue level. In a 50 µm mouse brain tissue, HCR-FISH targeting *ACTB*, IF using SMI-312 antibody (against neurofilament) and WGA stain (contrast adjusted to highlight blood vessels) were applied together. Image stacks were rendered in 3D and presented in the lower panel. The colors in images correspond to the following fluorescent dyes: blue – DAPI; green – Alexa488; gray – Alexa546; red – Alexa647. Scale bars (in pre-expansion units): 5 µm.

## Supplementary Tables

**Supplementary Table 1.**
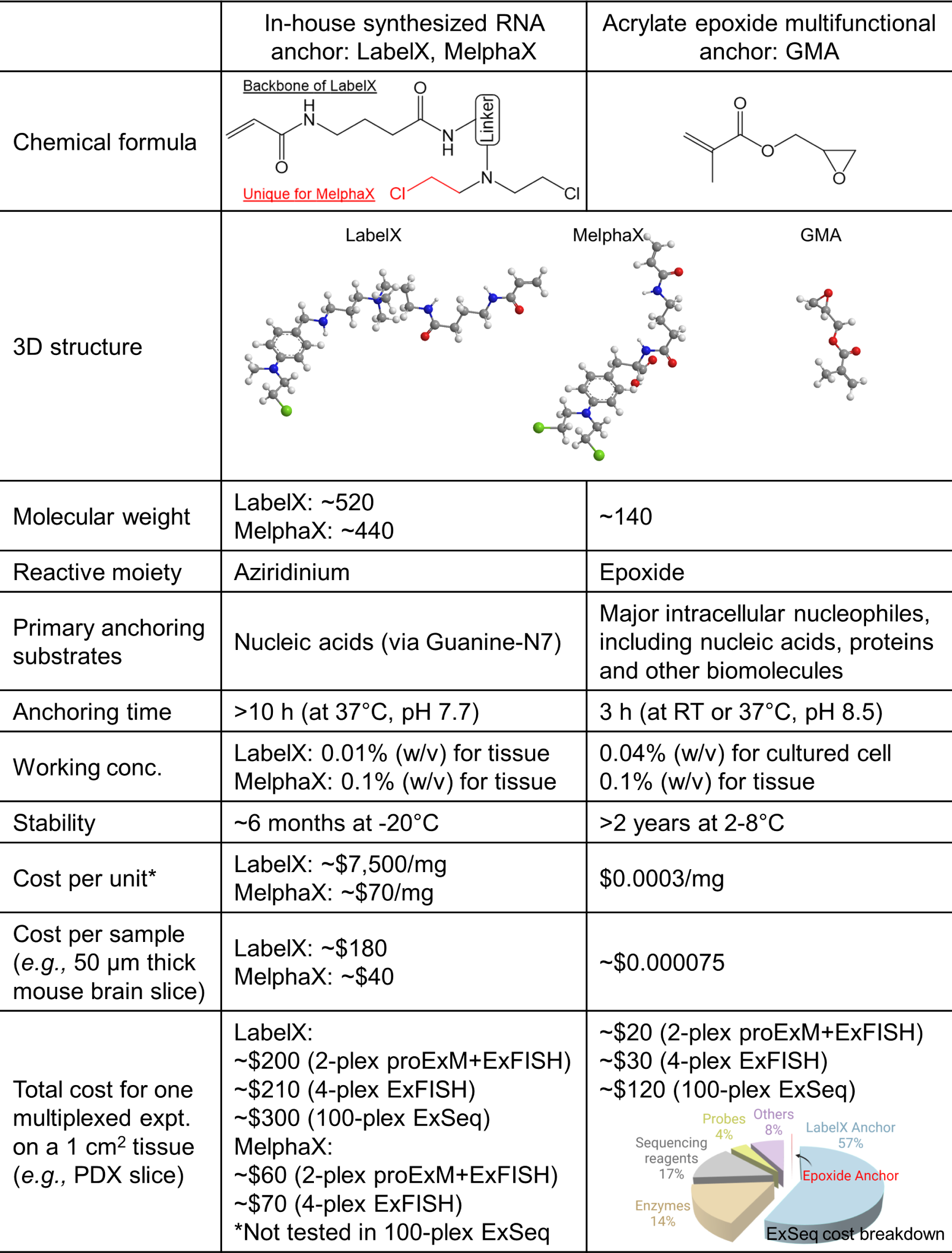
High-level comparison of LabelX, MelphaX and GMA for key parameters in multimodal ExM applications

**Supplementary Table 2.** Main antibodies used in this study

| **Antibody name** | **Vendor information** | **Catalog number** |
| --- | --- | --- |
| Mouse anti-tubulin | DSHB | E7 |
| Rabbit anti-GFP | ThermoFisher Scientific | A-6455 |
| Mouse anti-βII-spectrin | BD Biosciences | 612563 |
| Mouse anti-MAP2 | BioLegend | 801801 |
| Mouse anti-SMI312 | BioLegend | 837904 |
| Goat anti-mouse IgG Alexa 488 | ThermoFisher Scientific | A-21430 |
| Goat anti-rabbit IgG Alexa 555 | ThermoFisher Scientific | A-11017 |
| Goat anti-mouse IgG Alexa 546 | ThermoFisher Scientific | A-11018 |
| Goat anti-rabbit IgG Alexa 647 | ThermoFisher Scientific | A-21246 |

**Supplementary Table 3.** Main chemicals and reagents used in this study

| **Chemical/Reagent Name** | **Supplier** | **Catalog #** |
| --- | --- | --- |
| 16% Formaldehyde (w/v), methanol-free (PFA) | Thermo Fisher | 28908 |
| 4',6-diamidino-2-phenylindole (DAPI) | Sigma | D9542 |
| 4-Hydroxy-TEMPO (4-HT) | Sigma | 176141 |
| Acrylamide (AA) | Sigma | A9099 |
| Acrylamide/Bis 19:1, 40% (w/v) solution | Thermo Fisher | AM9022 |
| Acryloyl-X, SE, 6-((acryloyl)amino)hexanoic Acid, Succinimidyl Ester (AcX) | Thermo Fisher | A20770 |
| Aminoallyl-dUTP solution (50 mM) | Thermo Fisher | R1101 |
| Ammonium persulfate (APS) | Sigma | A3678 |
| Bind-silane | Sigma | GE17-1330-01 |
| BODIPY FL C<sub>12</sub> | Thermo Fisher | D3822 |
| CircLigase II ssDNA ligase | Lucigen | CL9025K |
| Deoxynucleotide (dNTP), 10 mM solution mix | NEB | N0447L |
| Dideoxynucleoside triphosphate set (ddNTP) | Roche | 03732738001 |
| DMEM | Thermo Fisher | 10569010 |
| DMSO, Anhydrous | Thermo Fisher | D12345 |
| DTT | Roche | 37016821 |
| DPBS, 1× | Corning | 21-031-CV |
| EDTA, 0.5M solution, pH 8.0 | Thermo Fisher | 15575020 |
| Endonuclease V | NEB | M0305S |
| Endoproteinase LysC | NEB | P8109S |
| Ethanolamine hydrochloride | Sigma | E6133 |
| Fetal bovine serum | Thermo Fisher | 16000036 |
| FM 1-43 FX membrane stain | Thermo Fisher | F35355 |
| Formamide (deionized) | Thermo Fisher | AM9344 |
| Glutaraldehyde (GA), 25% solution | Sigma | G5882 |
| Glycidyl methacrylate (GMA) | Sigma | 151238 |
| Guanidine hydrochloride | Sigma | 50937 |
| HCR-FISH probes, amplifiers and buffers | Molecular Instruments | N/A |
| Inosine | Sigma | I4125 |
| Label-IT amine | Mirus Bio | MIR3900 |
| MAXpack immunostaining media kit | Active Motif | 15251 |
| MES, 0.5M solution, pH 6.5 | Alfa Aesar | J63778 |
| MiSeq reagent kit v3 | Illumina | MS-102-3003 |
| N-(3-Dimethylaminopropyl)-N'-ethylcarbodiimide hydrochloride (EDC) | Sigma | 03450 |
| N,N,N',N'-Tetramethylethylenediamine (TEMED) | Sigma | T7024 |
| N,N'-Methylenebisacrylamide (BIS) | Sigma | M7279 |
| N-Hydroxysuccinimide (NHS) | Thermo Fisher | 24500 |
| Octadecyl rhodamine B chloride (R18) | Thermo Fisher | O246 |
| PBCV-1 DNA ligase | NEB | M0375L |
| PBS, 10× | Thermo Fisher | 70011044 |
| PEGylated bis(sulfosuccinimidyl)suberate (BE(PEG)9) | Thermo Fisher | 21582 |
| Penicillin-streptomycin (10,000 U/mL) | Thermo Fisher | 15140122 |
| phi29 DNA polymerase | Enzymatics | P7020-HC-L |
| Proteinase K | NEB | P8107S |
| RNase A | Thermo Fisher | EN0531 |
| RNase H | NEB | M0297L |
| RNase inhibitor | NEB | M0314S |
| Sigmacote | Sigma | SL2 |
| Sodium acrylate (SA) | Sigma | 408220 |
| Sodium bicarbonate, powder | Sigma | S6014 |
| Sodium borate, 0.5M solution, pH 8.5 | Alfa Aesar | J62902 |
| Sodium borohydride | Sigma | 213462 |
| Sodium chloride, 5M solution | Sigma | 59222C |
| Sodium dodecyl sulfate (SDS), 20% solution | Sigma | 05030 |
| SSC buffer, 20× | Promega | V4261 |
| Standard Taq reaction buffer (with magnesium chloride) | NEB | B9014S |
| SuperScript IV reverse transcriptase | Thermo Fisher | 18090050 |
| Terminal transferase | NEB | M0315L |
| TetraSpeck microspheres, 0.5 μm | Thermo Fisher | T7281 |
| Trimethylolpropane triglycidyl ether (TMPTE) | Sigma | 430269 |
| Tris buffer, 1M solution, pH 8.0, RNase-free | Thermo Fisher | AM9856 |
| Triton X-100 | Sigma | T8787 |
| Trypsin-EDTA (0.25% with phenol red) | Thermo Fisher | 25200072 |
| Tween 20 | Sigma | P9416 |
| UltraPure DNase/RNase-free distilled water | Thermo Fisher | 10977023 |
| Wheat germ agglutinin (WGA)-Alexa Fluor 647 | Thermo Fisher | W32466 |
| Zwittergent 3-10 detergent | Sigma | 693021 |

**Supplementary Table 4.** Anchoring conditions used in main experiments

| Figure | Sample type | Anchoring condition | Detection module |
| --- | --- | --- | --- |
| Fig. 1B (i) | Mouse brain slices | 0.1% GMA, 100 mM NaHCO<sub>3</sub> (pH 8.5), 3 h at 37°C | post-expansion HCR-FISH + YFP signal analysis |
| Fig. 1B (iii) | Human HeLa cells;<br>Mouse hippocampal neurons | 0.04% GMA, 100 mM NaHCO<sub>3</sub> (pH 8.5), 3 h at 37°C | pre-expansion IF + post-expansion HCR-FISH |
| Fig. 2A | Human HeLa cells | 0.04% GMA, 100 mM NaHCO<sub>3</sub> (pH 8.5), 3 h at RT | pre-expansion IF |
| Fig. 2B | Human HeLa cells | 0.04% GMA, 100 mM NaHCO<sub>3</sub> (pH 8.5), 3 h at RT | pre-expansion HCR-FISH + post-expansion HCR-FISH |
| Fig. 3 | Mouse hippocampal neurons | 0.04% GMA, 100 mM NaHCO<sub>3</sub> (pH 8.5), 3 h at 37°C | pre-expansion IF |
| Fig. 4B (ii)<br>Supp. Fig. 10A | Human HeLa cells | 0.04% GMA, 100 mM NaHCO<sub>3</sub> (pH 8.5), 6 h at RT | untargeted in situ sequencing |
| Fig. 4C | Human PDX breast cancer tissues | 0.1% GMA, 6 h at 4°C (with 1× PBS, pH 7.4) plus overnight at RT (with 100 mM NaHCO<sub>3</sub>, pH 8.5) | targeted in situ sequencing |
| Supp. Fig. 3 | Human HeLa cells;<br>Mouse brain slices | 0.04% GMA, 100 mM NaHCO<sub>3</sub> (pH 8.5), 3 h at RT;<br>0.1% GMA, 100 mM NaHCO<sub>3</sub> (pH 8.5), 3 h at 37°C (for mouse brain tissue) | post-expansion IF or post-expansion HCR-FISH |
| Supp. Fig. 4 | Human HeLa cells | 0.04% GMA, 100 mM NaHCO<sub>3</sub> (pH 8.5), 3 h at RT | DAPI staining before imaging |
| Supp. Fig. 5 | Human HeLa cells | as indicated in plots | post-expansion HCR-FISH |
| Supp. Fig. 6 | Human HeLa cells | 0.04% GMA, 100 mM NaHCO<sub>3</sub> (pH 8.5), 3 h at RT | post-expansion HCR-FISH |
| Supp. Fig. 7 | Mouse hippocampal neurons | 0.04% GMA, 100 mM NaHCO<sub>3</sub> (pH 8.5), 3 h at 37°C | pre-expansion IF |
| Supp. Fig. 8 | Mouse brain slices | 0.05% GMA + 0.05% TMPTE, 6 h at 4°C (with 1× PBS, pH 7.4) plus overnight at RT (with 100 mM NaHCO<sub>3</sub>, pH 8.5) | post-expansion YFP signal analysis |
| Supp. Fig. 10C | Mouse brain slices | 0.1% GMA, 6 h at 4°C (with 1× PBS, pH 7.4) plus overnight at RT (with 100 mM NaHCO<sub>3</sub>, pH 8.5) | targeted in situ sequencing |
| Supp. Fig. 11A-B | Human HeLa cells | 0.04% GMA, overnight at 4°C (with 1× PBS, pH 7.4) plus 3 h at RT (with 100 mM NaHCO<sub>3</sub>, pH 8.5) | pre-expansion or post-expansion lipid tag staining |
| Supp. Fig. 11C | Human HeLa cells | 0.04% GMA, 100 mM NaHCO<sub>3</sub> (pH 8.5), 3 h at RT | post-expansion WGA staining |
| Supp. Fig. 12A | Human HeLa cells | 0.04% GMA, overnight at 4°C (with 1× PBS, pH 7.4) plus 3 h at RT (with 100 mM NaHCO<sub>3</sub>, pH 8.5) | post-expansion R18 + WGA staining |
| Supp. Fig. 12B | Human HeLa cells | 0.04% GMA, overnight at 4°C (with 1× PBS, pH 7.4) plus 3 h at RT (with 100 mM NaHCO<sub>3</sub>, pH 8.5) | pre-expansion IF + post-expansion HCR-FISH + post-expansion R18 or WGA staining |
| Supp. Fig. 12C | Mouse brain slices | 0.1% GMA, overnight at 4°C (with 1× PBS, pH 7.4) plus 3 h at RT (with 100 mM NaHCO<sub>3</sub>, pH 8.5) | pre-expansion IF + post-expansion HCR-FISH + post-expansion WGA staining |

**Supplementary Table 5.** Gene list used for profiling SA501 PDX cancer tissues

| Proliferation | DNA repair | EMT | Immune Signaling | Metabolism | Migration | Transcription | Others |
| --- | --- | --- | --- | --- | --- | --- | --- |
| KRAS<br>SOX4<br>BCL2<br>CDK12<br>AKT1<br>NF1<br>ARHGDIB<br>PIK3CA<br>AKT3<br>SEPT4<br>IGF2R<br>AKT2<br>EGFR<br>BMP7 | BRCA1<br>PARP1<br>BRCA2<br>PARP3<br>RAD21<br>ATM<br>FANCL<br>ATR<br>POLE<br>FANCD<br>POLQ<br>RAD51D<br>RAD50 | VIM<br>SNAI2<br>TWIST<br>NOTCH2<br>NCAD<br>CD44<br>EPCAM<br>SNAI1 | HLA-C<br>B2M<br>CDKN2A<br>LGALS1<br>HLA-A | TXNL1<br>FBXO32<br>NFE2L2<br>SQLE<br>NDUFS5<br>CP<br>CRABP1<br>IDH2<br>SLC25A6<br>CTSV<br>HIF1A<br>COX5A<br>ALDH9A1<br>OAZ2<br>ANXA1<br>ARC<br>ATP5F1A<br>HACD3 | S100A11<br>CTNNB1<br>CCDC88A<br>SPARC<br>TMSB15A | RNPS1<br>RBP1<br>CAMTA1<br>ENY2<br>ZNF24<br>ELOB<br>WDR61<br>SNRNP25<br>FOXL2<br>DDX24<br>SNRPA1<br>CDCA7 | RPL17<br>RPS2<br>XIST<br>SEC11A<br>H1FO<br>UBE2I<br>HSPE1<br>COL5A1<br>MORF4L1<br>MESD<br>IER3IP1<br>PSMA4 |

